# Dysfunctional mechanotransduction regulates the progression of PIK3CA-driven vascular malformations

**DOI:** 10.1101/2024.08.22.609165

**Authors:** Wen Yih Aw, Aanya Sawhney, Mitesh Rathod, Chloe P. Whitworth, Elizabeth L. Doherty, Ethan Madden, Jingming Lu, Kaden Westphal, Ryan Stack, William J. Polacheck

## Abstract

Somatic activating mutations in *PIK3CA* are common drivers of vascular and lymphatic malformations. Despite common biophysical signatures of tissues susceptible to lesion formation, including compliant extracellular matrix and low rates of perfusion, lesions vary in clinical presentation from localized cystic dilatation to diffuse and infiltrative vascular dysplasia. The mechanisms driving the differences in disease severity and variability in clinical presentation and the role of the biophysical microenvironment in potentiating progression are poorly understood. Here, we investigate the role of hemodynamic forces and the biophysical microenvironment in the pathophysiology of vascular malformations, and we identify hemodynamic shear stress and defective endothelial cell mechanotransduction as key regulators of lesion progression. We found that constitutive PI3K activation impaired flow-mediated endothelial cell alignment and barrier function. We show that defective shear stress sensing in *PIK3CA^E542K^*endothelial cells is associated with reduced myosin light chain phosphorylation, junctional instability, and defective recruitment of vinculin to cell-cell junctions. Using 3D microfluidic models of the vasculature, we demonstrate that *PIK3CA^E542K^*microvessels apply reduced traction forces and are unaffected by flow interruption. We further found that draining transmural flow resulted in increased sprouting and invasion responses in *PIK3CA^E542K^* microvessels. Mechanistically, constitutive PI3K activation decreased cellular and nuclear elasticity resulting in defective cellular tensional homeostasis in endothelial cells which may underlie vascular dilation, tissue hyperplasia, and hypersprouting in *PIK3CA*-driven venous and lymphatic malformations. Together, these results suggest that defective nuclear mechanics, impaired cellular mechanotransduction, and maladaptive hemodynamic responses contribute to the development and progression of *PIK3CA*-driven vascular malformations.

## Introduction

Vascular malformations (VMs) are a class of rare genetic disorders associated with localized developmental abnormalities of venous, arterial, capillary, or lymphatic vessels [1–4]. Histologically, VMs are characterized by complexes of structural lesions with enlarged, irregular lumens that are lined with vascular endothelial cells (ECs) and surrounded by disorganized extracellular matrix [3, 5]. These lesions are often congenital, progress in severity over time, are associated with vascular obstruction and impaired drainage, and can be life threatening. Treatment for VMs, especially for complex malformations that occur in multiple tissues, is difficult and rarely curative, and the standard of care relies on surgery, sclerotherapy, and a limited repertoire of drugs.

VMs are caused by somatic mutations in genes that are involved in vascular development, and these same genes are commonly implicated in malignant tumor angiogenesis [4]. VMs are classified based on blood flow dynamics and are categorized as either slow-flow or fast-flow malformations [3]. The majority of slow-flow malformations, which do not involve an arterial component, are caused by mutations that result in upregulated phosphoinositide 3-kinase (PI3K) and mammalian target of rapamycin (mTOR) signaling [4]. In particular, somatic activating mutations of *PIK3CA* have been identified as the cause of the majority (∼80%) of cystic lymphatic malformations, and other pathologies, including generalized lymphatic anomalies, capillary malformations, and venous malformations [2, 6–9]. Several studies have demonstrated that overstimulation of PI3K signaling, independent of genetic mutations in *PIK3CA,* is associated with the pathological development of slow-flow VMs, including cerebral cavernous malformation [10–13] and fast-flow arteriovenous malformations in mouse models of hereditary hemorrhagic telangiectasia [14–16]. Given the implication of PI3K overactivation in both slow- and fast-flow VMs, understanding how hemodynamic forces contribute to the pathogenesis of *PIK3CA*-driven VMs could shed light on novel therapeutic approaches for various vascular anomalies.

*PIK3CA* encodes the p110α catalytic subunit (PI3Kα) of phosphatidylinositol-3-kinases (PI3K), a lipid kinase that is ubiquitously expressed and essential for regulating effector signaling pathways involved in cell proliferation, metabolism, migration, and survival [17]. At the cellular level, PI3K is activated by receptor tyrosine kinases, focal adhesion kinase, and G-protein coupled receptor signaling [17]. Upon activation, PI3K catalyzes the production of phosphatidylinositol 3,4,5-bisphosphate (PtdIns(3,4,5)P_3_ or PIP3) at the plasma membrane, leading to membrane translocation and downstream activation of AKT and mTOR signaling cascades that promote cell growth and survival [17]. PI3K signaling is also involved in the remodeling of the actin cytoskeleton. In particular, activation of PI3Kα has been demonstrated to modulate actomyosin contractility and cell motility in endothelial cells by regulating the activity of myosin light chain phosphatase [18], Rac1-[19], and RhoA-GTPases [20]. In addition to being regulated by growth factors signaling, PI3K functions downstream of the fluid shear stress sensitive mechanosensory complex that consists of platelet endothelial cell adhesion molecule 1 (PECAM-1), vascular endothelial (VE-) cadherin, and vascular endothelial growth factor receptor 2 (VEGFR2) [21–23]. While shear-stress induced activation of PI3K signaling is associated with endothelial cell elongation, alignment, and vasodilatory response, how overactivation of PI3K signaling impacts endothelial mechanotransduction remains unknown.

Multiple studies using mouse models have revealed that PI3Kα plays a crucial role in regulating venous and lymphatic vessel growth and development [6–9, 24–26]. Both the loss of function and inhibition of PI3Kα result in embryonic lethality due to deficient angiogenesis and venous fate specification [24, 27]. Conversely, expression of gain-of-function *PIK3CA^H1047R^* variant of PI3Kα in the endothelium leads to over-proliferation of venous and lymphatic endothelial cells [25, 26]. Interestingly, expression of the same oncogenic *PIK3CA^H1047R^* variant resulted in distinct vascular overgrowth and malformation defects in blood and lymphatic vessels [25]. Specifically, *PIK3CA-* driven vascular malformations manifest as locally restricted vessel overgrowth and dilation in veins and capillaries, but as dense hypersprouting vascular networks in the lymphatics [25]. However, the mechanisms by which one *PIK3CA* activating mutation produces distinct pathological phenotypes in different vascular beds remain to be investigated. The lack of arterial involvement in slow-flow VMs, together with common biophysical signatures of tissues susceptible to lesion formation including compliant extracellular matrix, low rates of perfusion, and drainage flow, suggest significant contributions of vascular hemodynamics and the biophysical microenvironment to *PIK3CA-*driven lesion progression.

Previously, we established a microphysiological model of VMs and identified that *PIK3CA* activating mutations result in elevated Rac1 activity in endothelial cells, which drives lesion formation through dysregulated cytoskeletal dynamics [19]. In this study, we address the role of fluid shear stress and the biophysical microenvironment in the development of *PIK3CA-*driven slow-flow malformations, and we identify hemodynamic shear stress and defective cellular mechanotransduction as key regulators of malformation progression. We demonstrate that endothelial cells expressing *PIK3CA^E542K^* activating mutation fail to elongate, align, and establish diffusive barrier function in response to laminar shear stress. We observe increased cellular and nuclear compliance, and we further identified deficient recruitment of vinculin, a mechanosensitive adaptor protein, to intercellular adherens junctions in human umbilical vein endothelial cells (HUVECs) expressing a PIK3CA activating mutation commonly associated with VMs. Additionally, using tissue-engineered microvessels, we demonstrate that mutant HUVECs form dilated microvessels that are insensitive to flow interruption and are hypersensitive to basal-to-apical transmural drainage, resulting in sprouting and invasion responses to transmural flow. These results demonstrate that altered endothelial cellular and nuclear mechanical properties, as well as dysfunctional mechanotransduction of hemodynamic stresses, contribute to pathological vascular overgrowth and hypersprouting in *PIK3CA*-driven VMs.

## Results

### Constitutive PI3K activation impedes flow-mediated ECs remodeling and alignment to laminar shear stress

The development of PIK3CA-driven vascular lesions is restricted to slow-flow regions of the vasculature [28]. To determine whether VM causal mutations in *PIK3CA* impact mechanotransduction of hemodynamic shear stress, we generated human umbilical vein endothelial cells (HUVECs) expressing wild-type *PIK3CA* (*PIK3CA^WT^*) or *PIK3CA* harboring a *PI3K-*activating mutation (*PIK3CA^E542K^*), as described previously [19], and applied flow via orbital shaking (**Figure 1**). This approach allowed us to apply a range of shear stresses, both laminar and oscillatory, to cultured *PIK3CA^WT^*and *PIK3CA^E542K^* monolayers. At a wall shear stress magnitude similar to that experienced by venous ECs (∼11 dyn/cm^2^) [29, 30] near the edge of the well, *PIK3CA^WT^* HUVECs became highly elongated and aligned with the local flow direction (**Figure 1A-C**). In response to lower magnitude wall shear stress and oscillatory flow toward the center of the well, *PIK3CA^WT^* HUVECs adopted a more isometric morphology (**Figure 1A, C**). In contrast, *PIK3CA^E542K^* HUVECs remained isometric at all magnitudes of wall shear stress (**Figure 1A-C**). We further observed increased cell density in *PIK3CA^E542K^* HUVECs across all applied shear stress magnitudes, indicating impaired flow-mediated quiescence with excess PI3K activation (**Figure 1D**). Consistent with orbital shaking results, *PIK3CA^E542K^* HUVECs remained misaligned and overcrowded in response to 10 dyn/cm^2^ laminar shear stress applied in a Hele-Shaw flow cell (**Figure 1F**).

**Figure 1.**
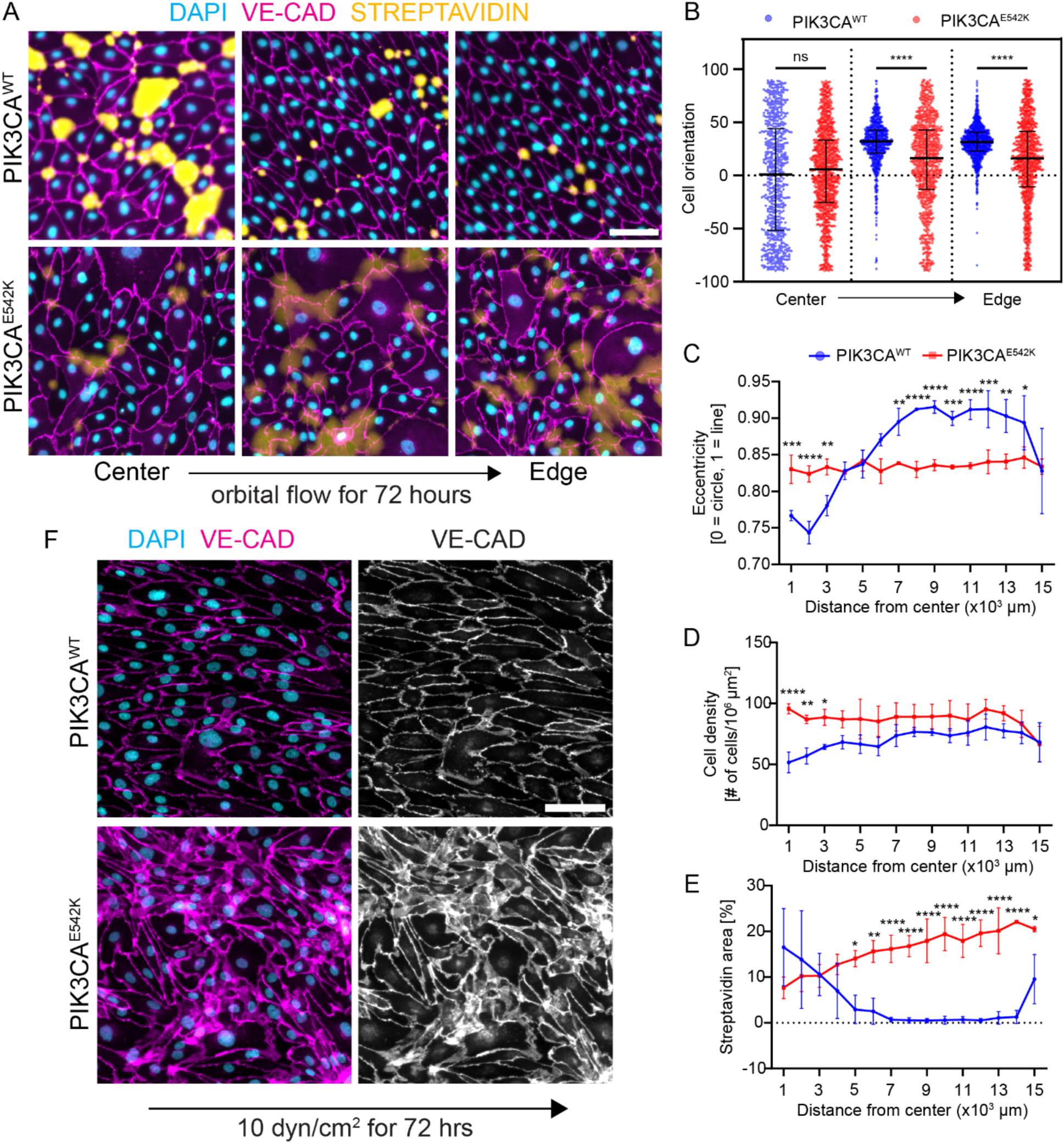
Constitutive PI3K activation impedes shear stress-induced endothelial cell alignment and vascular barrier function. (**A**) Representative images of *PIK3CA^WT^*and *PIK3CA^E542K^* HUVECs cultured on biotinylated fibronectin for 72 hr under orbital shaking conditions. Cells were labeled with DAPI (cyan) and VE-cadherin (magenta). Monolayer permeability was assessed through streptavidin labeling (yellow). Scale bar = 10 µm. (**B**) Quantification of cellular orientation in the central, middle, or outer edge region of the well after exposure to orbital shaking (n = 3; mean ± s.d., Kruskal-Wallis test followed by Dunn’s test). (**C**) Quantification of cellular eccentricity as a function of radial distance from the center of the well for *PIK3CA^WT^* and *PIK3CA^E542K^* HUVECs (n = 3; mean ± s.d., two-way analysis of variance (ANOVA) followed by Šidák’s test). (**D**) Quantification of cell density as a function of radial distance from the center of the well for PIK3CA^WT^ and PIK3CA^E542K^ HUVEC (n = 3; means ± s.d., two-way ANOVA followed by Šidák’s test). (**E**) Quantification of streptavidin intensity as a function of radial distance from the center of the well for *PIK3CA^WT^* and *PIK3CA^E542K^* HUVECs (n = 3; mean ± s.d., two-way ANOVA followed by Šidák’s test). (**F**) Representative images of *PIK3CA^WT^* and *PIK3CA^E542K^*HUVECs exposed to 10 dyn/cm^2^ laminar shear stress in a Hele-Shaw flow cell. Scale bar = 10 µm. *p< 0.05, **p< 0.01, ***p< 0.001, ****p< 0.0001 for all plots.

To determine if the defective mechanotransduction we observed in blood endothelial cells could play a role in lymphatic malformation progression, we generated lymphatic endothelial cells (LECs) expressing either *PIK3CA^WT^* or *PIK3CA^E542K^* (**Figure S1**). Similar to *PIK3CA^E542K^* HUVECs, we observed that *PIK3CA^E542K^* LECs were larger in cell size and more isometric compared to *PIK3CA^WT^* LECs in response to orbital shaking (**Figure S1**), suggesting that excess PI3K activation and effector signaling perturb mechanotransduction of hemodynamic shear stress in both the blood and the lymphatic endothelium. To determine the functional consequences of cellular misalignment to flow, we measured the permeability of *PIK3CA^WT^*and *PIK3CA^E542K^* HUVEC monolayers in response to flow. We cultured *PIK3CA^WT^* and *PIK3CA^E542K^* HUVEC under orbital shaking conditions on biotinylated-fibronectin substrates, and permeability was quantified by fluorescence intensity of Cy3-conjugated streptavidin that bound to the biotinylated surface (**Figure 1A, E**). In agreement with previous reports [31], we observed improved barrier function in *PIK3CA^WT^*HUVECs at regions corresponding to high laminar shear stress and a gradual increase in permeability with decreasing wall shear stress magnitude (**Figure 1A, E**). Interestingly, *PIK3CA^E542K^* HUVECs demonstrated a dichotomous relationship between shear stress magnitude and barrier function, with increased permeability as compared to wild-type cells at high magnitudes of shear stress and reduced permeability at lower shear stress magnitudes (**Figure 1A, E**). Collectively, these findings demonstrated that constitutive PI3K activation hinders flow-mediated endothelial cell alignment and barrier function.

### Priming with hemodynamic shear stress prior to *PIK3CA^E542K^* induction restores cell alignment but does not impact permeability

To establish the temporal relationship between elevated PI3K signaling, defective endothelial cell remodeling, and loss of vascular barrier function, we generated HUVECs expressing either *tet-ON-PIK3CA^WT^* or *tet-ON-PIK3CA^E542K^* in which the expression of WT or mutant variant of *PIK3CA* can be induced by the addition of doxycycline (**Figure S2A**). Treatment with doxycycline to induce *PIK3CA^WT^ or PIK3CA^E542K^* expression induced similar levels of p110⍺ expression as measured by Western blot, indicating a comparable level of lentivirus integration and protein overexpression in both cell lines (**Figure S2A**). Upon addition of doxycycline, we observed increased levels of phospo-AKT1(Ser473) in *tet-ON-PIK3CA^E542K^*, but not in *tet-ON-PIK3CA^WT^* endothelial cells, demonstrating activation of PI3K effector signaling in response to induction (**Figure S2A**). To verify the functional relevance of this construct, we generated 3D human microvascular networks with *tet-ON-PIK3CA^E542K^* HUVECs (**Figure S2B-C**). We observed that the addition of doxycycline during seeding of *tet-ON-PIK3CA^E542K^* HUVECs accelerated lumenogenesis and resulted in the formation of microvascular networks with enlarged and irregular lumens (**Figure S2B**), in agreement with our previous study demonstrating that constitutive activation of PIK3CA^E542K^ resulted in pathologic network formation [19].

To establish the temporal relationships between dysregulated PI3K activation and mechanotransduction of hemodynamic shear stress, we induced expression of *PIK3CA^E542K^* either prior to or after the application of flow (**Figure 2A**). When *PIK3CA^E542K^* expression was induced before the application of flow, ECs remained unresponsive to shear stress, as measured by cell orientation and elongation (**Figure 2A-C**). However, when *PIK3CA^E542K^* expression was triggered one day after flow, flow-conditioned HUVECs remained elongated and aligned parallel to the flow direction (**Figure 2A-C**). Interestingly, although flow preconditioning rescued cell morphology and alignment, PI3K activation significantly reduced vascular barrier function compared with uninduced (no doxycycline) control (**Figure 2D**). Collectively, these results indicate a mechanistic separation between flow-mediated cell alignment and barrier function, whereby excessive PI3K signaling may have a more direct and immediate impact on EC adherens junction stability and resultant permeability.

**Figure 2.**
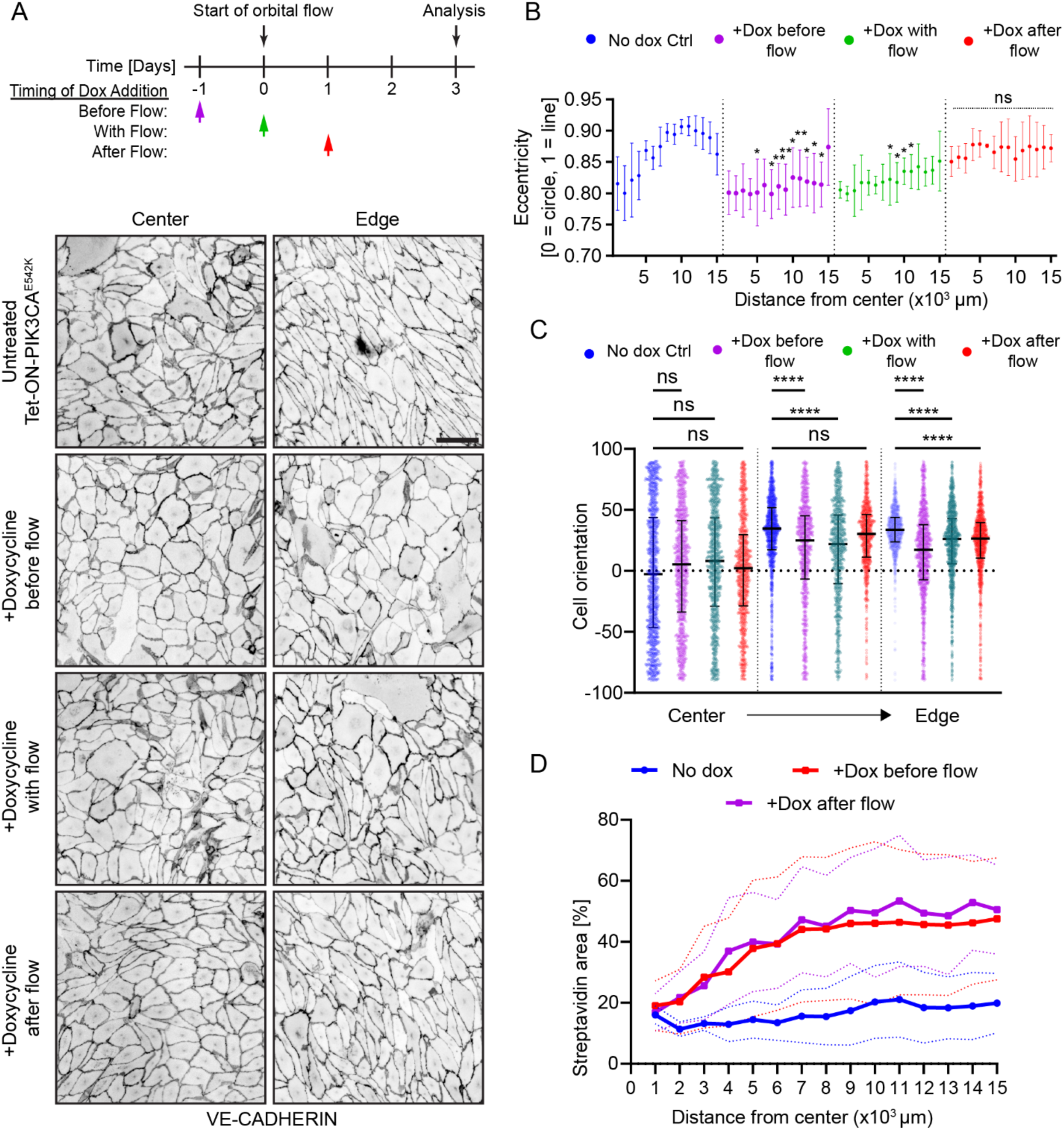
Preconditioning with shear stress prior to the induction of *PIK3CA^E542K^* expression restores shear stress-mediated EC alignment but not barrier function. (**A**) Schematic of experimental workflow and representative images of VE-cadherin staining in control cells and cells where expression of *PIK3CA^E542K^* was induced 24 hr before flow, at the onset of flow, or 24 hr after flow. Images were captured near the center of the well or near the outer edge of the well. Scale bar = 10 µm. (**B**) Quantification of cellular eccentricity as a function of radial distance for control cells as well as for cells induced with doxycycline 24 hr before flow, at the onset of flow, or 24 hr after flow (n = 3; mean ± s.d., two-way ANOVA followed by Šidák’s test). (**C**) Quantification of cellular orientation in the central, middle, or outer region of the well for cells induced with doxycycline 24 hr before flow, at the onset of flow, or 24 hr after flow (n = 3; mean ± s.d., Kruskal-Wallis tests followed by Dunn’s test). (**D**) Quantification of streptavidin area as a function of radial distance from the center of the well for control cells, cells induced with doxycycline 24 hr before flow, and cells induced with doxycycline 24 hr after flow (n = 3; mean ± s.d. (dotted lines), ANOVA followed by Dunnett test). *p< 0.05, **p< 0.01, ***p< 0.001, ****p< 0.0001 for all plots.

### Mutant ECs are characterized by adherens junction instability

We previously reported the presence of discontinuous VE-cadherin-containing adherens junctions, decreased cortical actin localization, and increased stress fibers formation in HUVECs expressing *PIK3CA* activating mutations [19]. We observed similar decreased in the distribution of cortical actin, lack of stress fibers alignment, and reduced VE-cadherin signal in cell-cell contacts of flow-conditioned *PIK3CA^E542K^* HUVECs (**Figure 3A-D**). Additionally, we observed increased microtubule density and similar loss of microtubule and vimentin alignment in *PIK3CA^E542K^* HUVECs (**Figure S3 and S4**). These findings suggest that the mechanical coupling between the cytoskeleton and adhesion complexes and stability of VE-cadherin adhesion complexes may be hampered with constitutive PI3K activation. To quantify the stability of VE-cadherin intercellular adherens junctions, a VE-cadherin pulse-chase experiment was performed (**Figure 4A-C**). Surface-pool of VE-cadherin were pulse-labeled with non-functional blocking antibody and turnover of labeled VE-cadherin complexes were quantified after a 2-hour chase period. We observed a reduced level of labeled VE-cadherin at cell-cell contacts and a significant increase in the number of internalized labeled VE-cadherin positive vesicles within the cytoplasm of *PIK3CA^E542K^* HUVECs (**Figure 4A-C**), indicating that adherens junctions are indeed less stable in mutant endothelium. Given that endothelial mechanotransduction of shear stress is partially mediated by cortical Src kinase activation driven by the formation of junctional VE-cadherin/PECAM-1/VEGFR2 mechanosensory complex [23], we investigated the dynamics of Src kinase activation in *PIK3CA* mutant endothelium (**Figure 4D-E**). Interestingly, we found that the temporal dynamics of Src activation measured by the level of phospho-Src (Tyr416) was similar both in magnitude and temporal activation among untransduced HUVEC, PIK3CA^WT^, and PIK3CA^E542K^ HUVECs (**Figure 4D-E**), suggesting that while assembly and turnover of adherens junctions are disrupted in mutant endothelial cells, some elements of downstream effector signaling remain functional.

**Figure 3.**
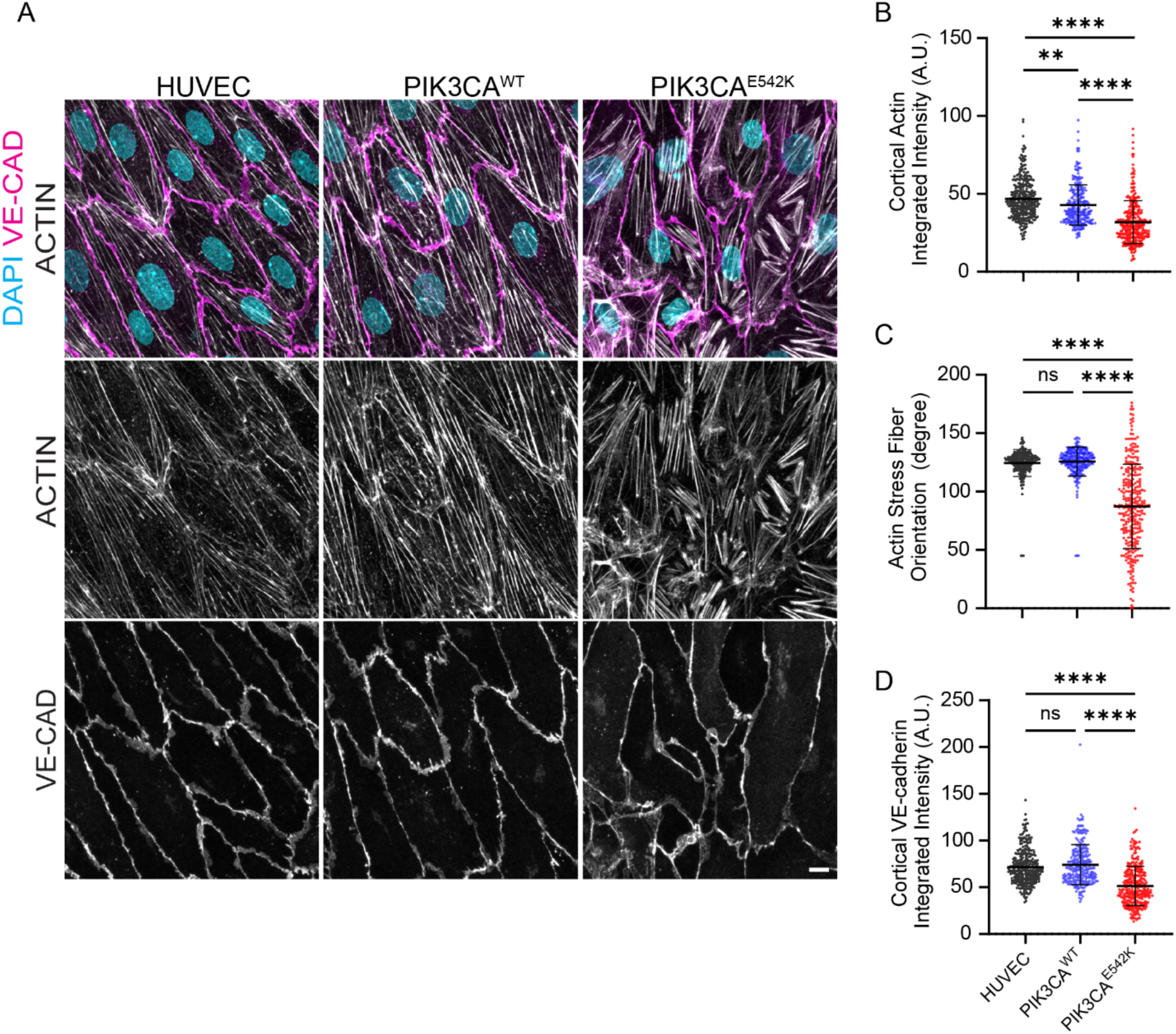
Loss of stress fiber alignment, reduced cortical actin formation, and decreased VE-cadherin accumulation at cell-cell contacts in *PIK3CA^E542K^* endothelium. (**A**) Representative confocal images of untransduced HUVECs, *PIK3CA^WT^*, and *PIK3CA^E542K^* HUVECs stained for DAPI (blue), actin (grey), and VE-cadherin (magenta). Cells were cultured under orbital flow for 72 hours. Scale bar = 10 µm. (**B**) Quantification of the integrated intensity of actin fluorescence signal at cell-cell contacts (n = 320 untransduced HUVECs, 244 *PIK3A^WT^* and 297 *PIK3CA^E542K^* cells; mean ± s.d., one-way ANOVA followed by Tukey’s test). (**C**) Angular distribution of stress fiber orientation in flow-conditioned untransduced HUVECs, *PIK3CA^WT^*, and *PIK3CA^E542K^* HUVECs (n = 324 untransduced HUVECs, 248 *PIK3A^WT^* and 296 *PIK3CA^E542K^* cells; mean ± s.d., one-way ANOVA followed by Kruskal-Wallis test). (**D**) Quantification of endogenous VE-cadherin fluorescence signal at cell-cell contacts (n = 320 untransduced HUVECs, 244 *PIK3A^WT^* and 297 *PIK3CA^E542K^* cells; mean ± s.d., one-way ANOVA followed by Tukey’s test).

**Figure 4.**
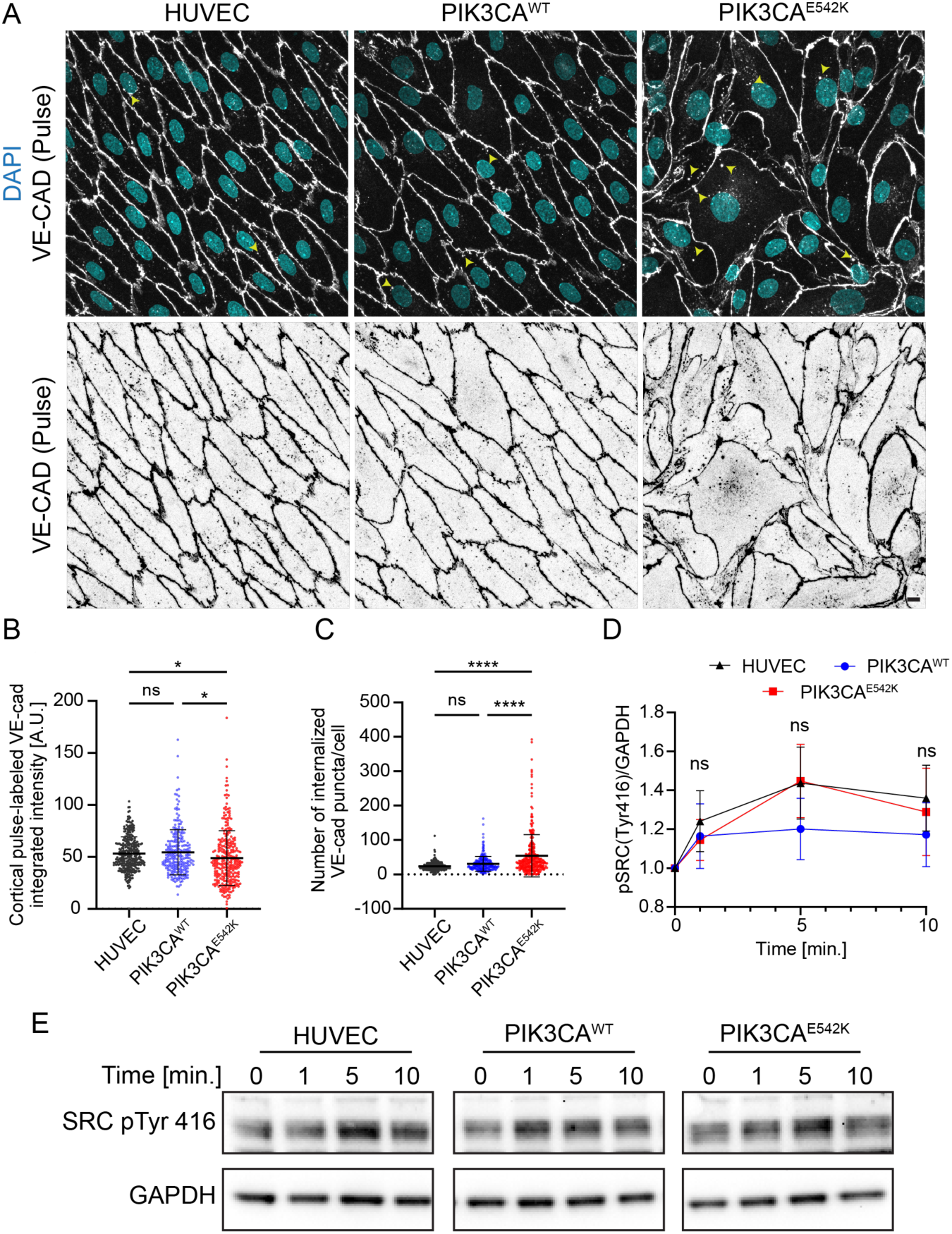
Increased junctional VE-cadherin turnover with normal shear-stress induced SRC phosphorylation in mutant *PIK3CA^E542K^* ECs. (**A**) Representative confocal images of untransduced HUVECs, *PIK3CA^WT^*, and *PIK3CA^E542K^* HUVECs, where the surface-pool of VE-cadherin (grey) was pulse-labeled with a fluorescently conjugated VE-cadherin antibody prior to fixation and permeabilization. Unbound antibodies were washed, and cells were fixed and stained with DAPI (blue) after a 2-hour chase period with media. Yellow arrowheads indicate internalized pulse-labeled VE-cadherin puncta. Scale bar = 10 µm. (**B**) Quantification of pulse-labeled VE-cadherin intensity at cell-cell contacts.junctions. (**C**) Quantification of the number of VE-cadherin-positive puncta per cell (n = 316 untransduced HUVECs, 245 *PIK3A^WT^*and 294 *PIK3CA^E542K^* cells; mean ± s.d., one-way ANOVA followed by Tukey’s test). (**D**) Changes in phospho-SRC(Y416) (active-Src) level in untransduced HUVECs, *PIK3CA^WT^*, and *PIK3CA^E542K^*HUVECs in response to the application orbital flow (n = 3; mean ± s.d., two-way ANOVA followed by Tukey’s test). (**E**) Representative western blot analysis of phospho-SRC(Y416) levels in HUVECs, *PIK3CA^WT^*, and *PIK3CA^E542K^* HUVECs subjected to laminar shear stress through orbital shaking for the indicated time points.

### Defective recruitment of vinculin at adherens junctions in PIK3CA mutant endothelium

VE-cadherin containing junctions are mechanically connected to the actin cytoskeleton through the interaction of VE-cadherin with β- and α-catenin [32, 33]. Under mechanical load, tensile stress unfolds α-catenin, allowing for vinculin binding to α-catenin and subsequent force-dependent strengthening of cell-cell junctions [34–36]. To address whether the mechanical coupling between actin cytoskeleton and adhesion complexes could underlie the increased permeability in mutant endothelium, we investigated the spatial localization of vinculin in *PIK3CA^WT^* and *PIK3CA^E542K^* HUVECs (**Figure 5A-C**). In *PIK3CA^WT^* HUVECs, vinculin localized at the cell cortex and co-localized with VE-cadherin containing cell-cell adhesions and along actin stress fibers (**Figure 5A-B**). In contrast, *PIK3CA^E542K^* HUVECs demonstrated reduced recruitment of vinculin to intercellular adherens junctions as compared to *PIK3CA^WT^* HUVECs (**Figure 5A-B**). We further observed that while vinculin primarily localizes in a straight and linear pattern in focal adhesion plaques of *PIK3CA^WT^*HUVECs, vinculin appears as larger, dot-like structures in flow-conditioned *PIK3CA^E542K^* HUVECs (**Figure 5C**). Despite significant changes in vinculin localization, western blot analysis showed no measurable differences in the amounts of VE-cadherin, β-catenin, and vinculin between control and mutant HUVECs (**Figure 5D-E**). Interestingly, in probing for changes for overall actomyosin contractility, we observed reduced level of phosphorylated myosin light chain II (pMLC2) in mutant ECs compared with control ECs (**Figure 5D-E**). These data demonstrate defective transmission of mechanical forces and stabilization of cell-cell and cell-matrix adhesion complexes in *PIK3CA^E542K^*HUVECs.

**Figure 5.**
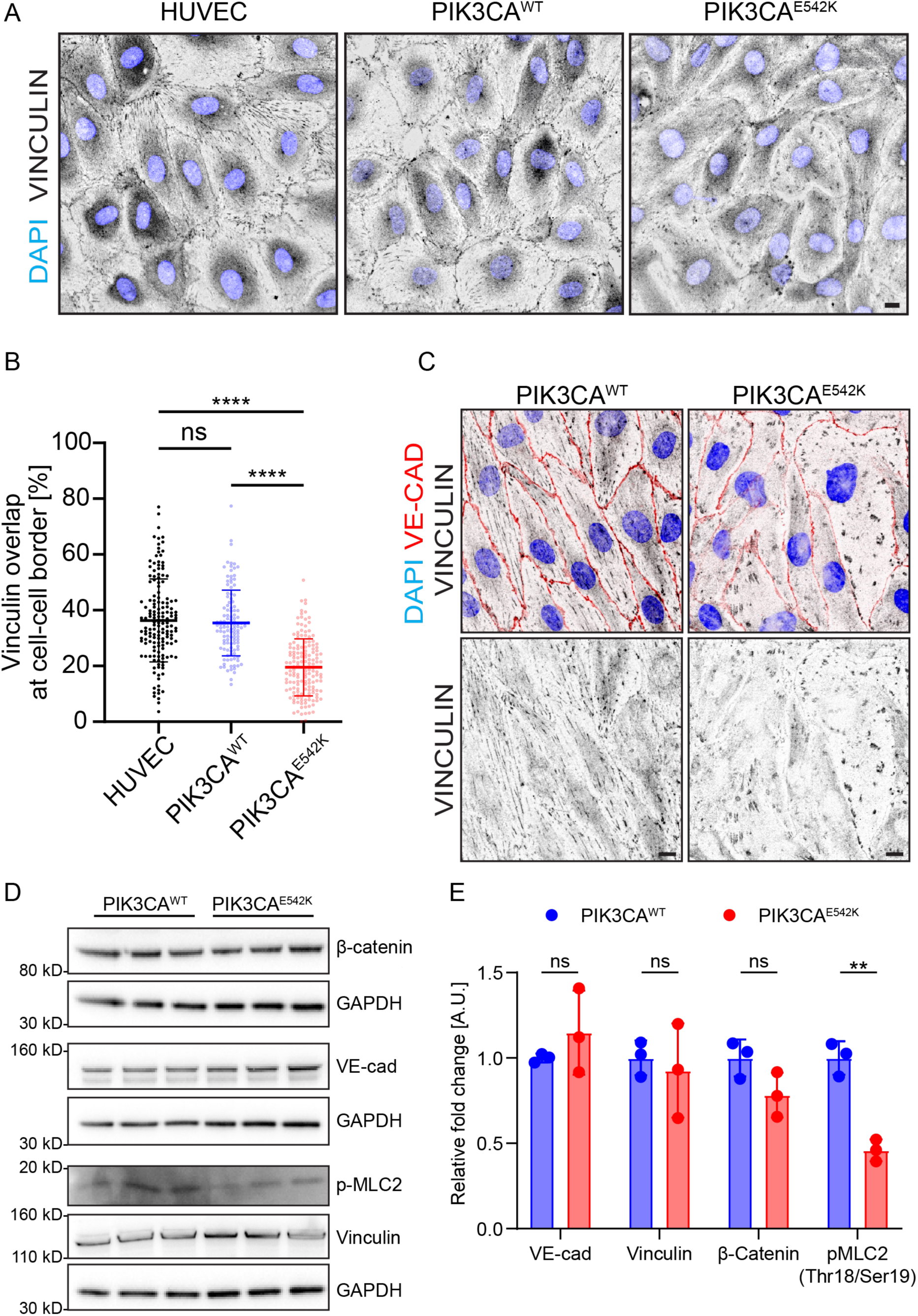
Loss of junctional vinculin recruitment in *PIK3CA^E542K^* endothelium. (**A**) Representative confocal images of untransduced HUVECs, *PIK3CA^WT^* and *PIK3CA^E542K^* HUVECs stained for vinculin (grey), and DAPI (blue). Scale bar = 10 µm. (**B**) Quantification of percent vinculin area at cell-cell contacts (n = 119 untransduced HUVECs, 157 *PIK3A^WT^* and 156 *PIK3CA^E542K^*cells; mean ± s.d., one-way ANOVA followed by Tukey’s test). (**C**) Representative confocal images of vinculin localized at focal adhesions in flow-conditioned *PIK3CA^WT^* and *PIK3CA^E542K^*. Cells were partially permeabilized during fixation and stained for DAPI (blue) VE-cadherin (red) and vinculin (grey). Scale bar = 10 µm. (**D**) Western blots and (**E**) quantification for β-catenin, VE-cadherin, phospho-myosin light chain 2, and vinculin level in *PIK3CA^WT^*or *PIK3CA^E542K^* cells (n = 3; mean ± s.d., two-tailed unpaired t-test). *p< 0.05, **p< 0.01, ***p< 0.001, ****p< 0.0001 for all plots.

### PIK3CA^E542K^ HUVECs form dilated microvessels with increased sprouting and reduced traction forces

To relate our findings in 2D cell culture to the pathophysiological phenotype of vascular malformations, we generated and characterized the effect of flow on *PIK3CA^WT^* or *PIK3CA^E542K^*tissue engineered human microvessels (**Figure 6**). Engineered microvessels were molded using an acupuncture needle to generate a hollow 160-μm cylindrical channel lined with HUVECs and fully embedded within 2.5 mg/mL type I collagen hydrogel [31, 37]. Cyclic flow of approximately 1-3 dyn/cm^2^ was imparted by culturing microvessels on a laboratory rocker [37]. After two days of culture under a slow flow environment, *PIK3CA^WT^*HUVECs formed uniform microvessels with a limited number of sprouts and no apparent HUVEC invasion into the subluminal matrix (**Figure 6A-B**). In contrast, *PIK3CA^E542K^* microvessels were significantly dilated and developed numerous sprouts along the vessel length (**Figure 6A-B**). The expanded mutant vessel diameter coincides with an increased number of *PIK3CA^E542K^*HUVECs, implicating defective regulation of flow-mediated quiescence with excessive PI3K signaling. Treatment with rapamycin, an mTOR inhibitor, or alpelisib, a PIK3CA-specific kinase inhibitor, reduced vessel dilation (**Figure 6A-B**). Importantly, when compared to rapamycin, we observed a more effective reduction in number of sprouts with alpelisib treatment (**Figure 6A-B**). This observation is consistent with our previous finding and the notion that in addition to contributing to uncontrolled growth through downstream mTORC1/2 and growth signaling activation, PI3K hyperactivation alters cytoskeletal dynamics and mechanical properties of endothelial cells [19].

**Figure 6.**
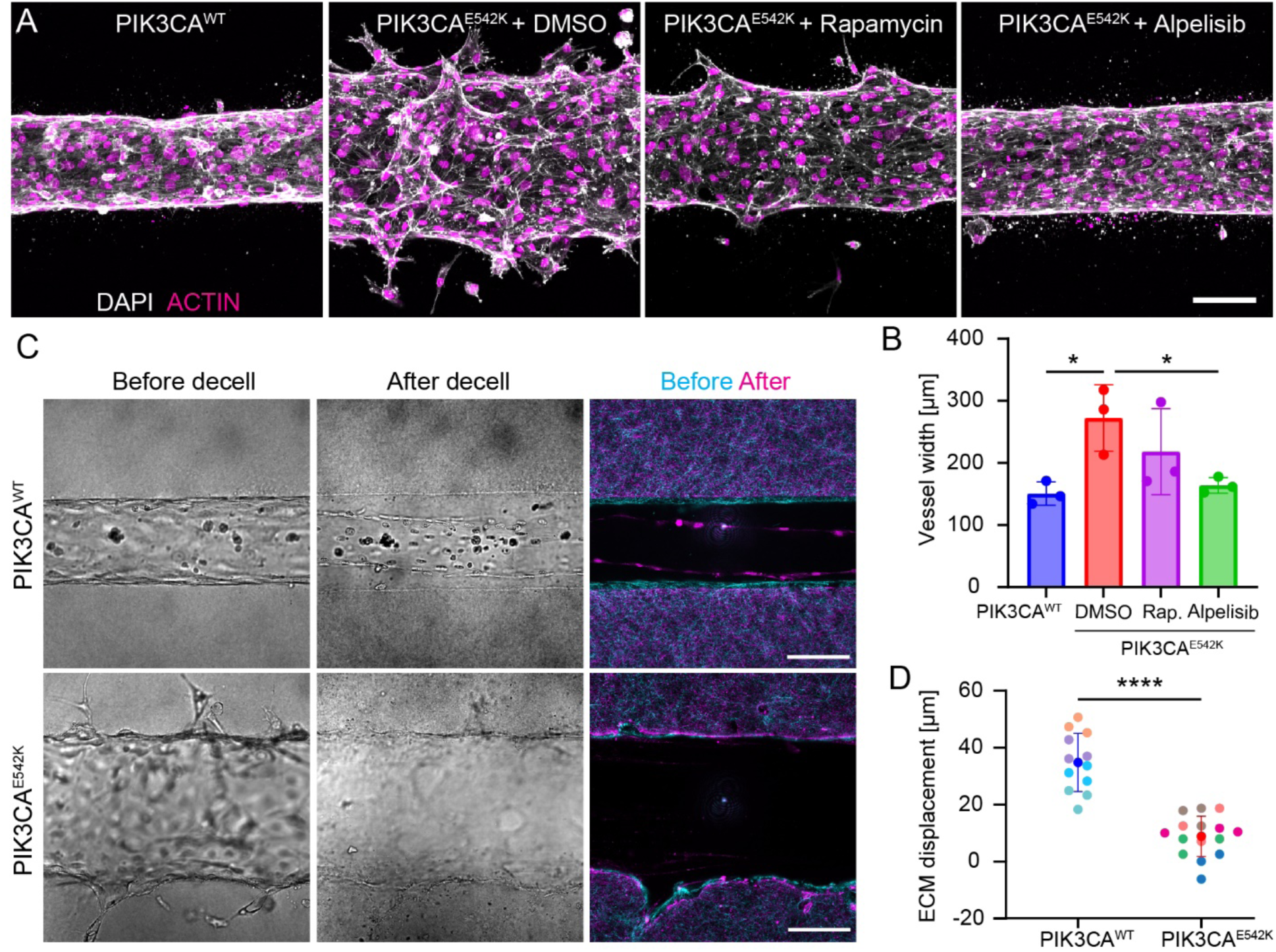
PIK3CA^E542K^ ECs form dilated microvessels with increased sprouting and reduced traction forces. (**A**) Representative confocal images (maximum intensity projections) of *PIK3CA^WT^* and *PIK3CA^E542K^* microvessels treated with DMSO, 1 µM alpelisib, or 1 µM rapamycin. Scale bar = 100 µm. (**B**) Quantification of vessel width in *PIK3CA^WT^* microvessels or *PIK3CA^E542K^*microvessels treated with DMSO load control, rapamycin (rap.), or alpelisib (n = 3; mean ± s.d., one-way ANOVA followed by Tukey test). (**C**) Representative phase contrast (grey) and confocal reflectance (cyan and magenta) images of *PIK3CA^WT^* and *PIK3CA^E542K^* microvessels before and after decellularization. (**D**) Quantification of subluminal ECM displacement before and after decellularization of *PIK3CA^WT^* or *PIK3CA^E542K^* microvessels (n ≥ 4 microvessels, 3 measurements per microvessel; data points from same devices are color-matched; mean ± s.d., two-tailed unpaired t-test). *p< 0.05, **p< 0.01, ***p< 0.001, ****p< 0.0001 for all plots.

Given the defective recruitment of vinculin to adherens junctions, increased vascular permeability, loss of cortical actin organization, and prominent stress fiber formation in *PIK3CA^E542K^*HUVECs, we reasoned that the dilated microvessel phenotype may reflect reduced cell-cell adhesion strength and altered cell-matrix traction forces in mutant microvessels. To assess changes in cell-generated forces, we visualized the collagen matrix surrounding the microvessels through reflectance imaging and compared the displacement of matrix before and after lysing ECs with detergent (**Figure 6C-D**). Notably, we observed significantly lower matrix displacement in mutant microvessels when compared to control (**Figure 6B-D**), indicating reduced traction forces in *PIK3CA^E542K^*microvessels.

### PIK3CA^E542K^ microvessels are unresponsive to flow interruption

To determine whether the slow-flow environment of native lesions potentiates VM progression, we next investigated the effect of transitioning from flow to static culture of *PIK3CA^WT^* and *PIK3CA^E542K^* microvessels by preconditioning microvessels with flow for 24 hrs then transitioning to static conditions for the subsequent 24 hrs. After transition to static culture, *PIK3CA^WT^* HUVECs delaminated and detached into the apical lumen, resulting in gaps in the endothelium (**Figure 7A**). In contrast, *PIK3CA^E542K^* HUVECs were insensitive to flow interruption, with no observable delamination and the formation of tortuous, sprouted, and dilated microvessels that are akin to mutant vessels cultured under flow conditions (**Figure 7B-C**). In all culture conditions, *PIK3CA^E542K^* microvessels contained approximately twice the number of cells and were approximately 2-fold wider than *PIK3CA^WT^* microvessels (**Figure 7C**). While *PIK3CA^E542K^*microvessels showed minimal effects from flow interruption, flow interruption in *PIK3CA^WT^* microvessels resulted in gaps covering approximately 6.5% of the vessel area (**Figure 7C**). Similar lack of cell delamination and cellular invasion were observed in *PIK3CA^E542K^* microvessels cultured under fully static condition (**Figure S5A-C**). Collectively, these findings indicate that constitutive PI3K activation renders ECs insensitive to vascular regression upon flow interruption and may lead to improved survival and favorable expansion of mutant ECs in the no flow and slow flow characteristics of PIK3CA-driven VMs.

**Figure 7.**
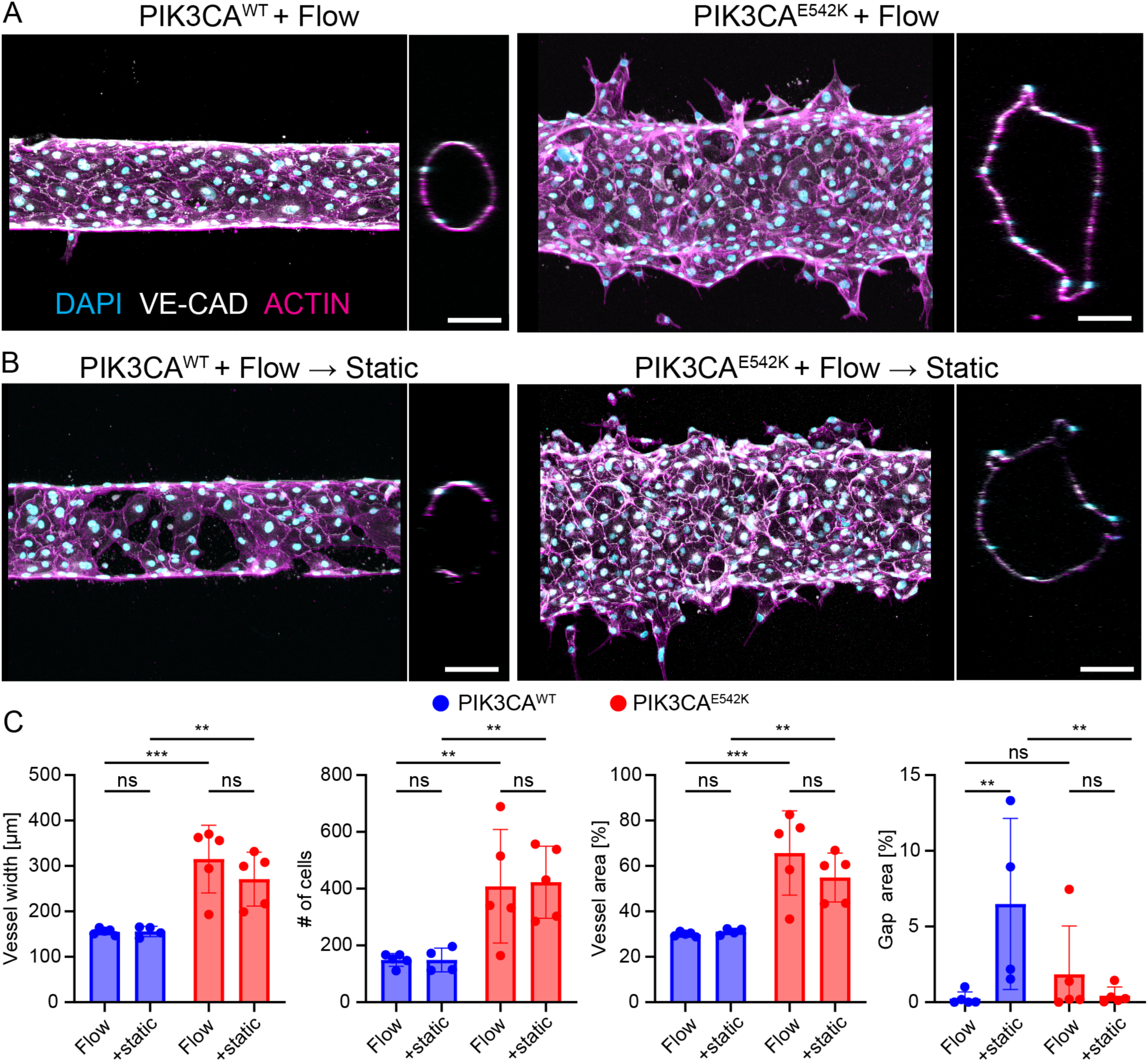
*PIK3CA^E542K^* microvessels are unresponsive to flow interruption. (**A**) Representative images of microvessels generated from *PIK3CA^WT^* and *PIK3CA^E542K^* HUVECs. Microvessels were exposed to continuous rocking 48 hr and stained for DAPI (cyan), VE-cadherin (white), and F-actin (magenta). Scale bars = 100 µm. (**B**) Representative images of *PIK3CA^WT^* and *PIK3CA^E542K^*microvessels exposed to 24 hr rocking followed by 24 hr static conditions and stained for DAPI (cyan), VE-cadherin (white), and F-actin (magenta). Scale bars = 100 µm. (**C**) Quantification of vessel width, number of cells, vessel area, and gap area for *PIK3CA^WT^* and *PIK3CA^E542K^*microvessels exposed to either continuous or interrupted flow (n ≥ 4; mean ± s.d., two-way ANOVA). *p< 0.05, **p< 0.01, ***p< 0.001, ****p< 0.0001 for all plots.

### Basal-to-apical transmural flow exacerbates hypersprouting in PIK3CA mutant microvessels

In addition to luminal shear stress, venous and lymphatic ECs are subjected to draining transmural flow. Hydrostatic and oncotic pressure differences across the vessel wall causes fluid to extravasate from blood capillaries, resulting in the accumulation of interstitial fluid in the surrounding matrix which is subsequently reabsorbed and collected by venous and lymphatic vessels [38]. Therefore, in addition to experiencing low level of wall shear stress, venous and lymphatic ECs are also subjected to fluid stresses arising from transmural drainage. To investigate the effect of transmural flow on control HUVECs, *PIK3CA^WT^*, and *PIK3CA^E542K^* microvessels, we cultured control or mutant HUVECs microfluidic devices that contain two parallel microvessels in a shared collagen I hydrogel [39]. This two-channel configuration allows us to establish a pressure gradient between the two microchannels to drive transmural flow into (draining vessel, basal-to-apical flow) or out of (source vessel, apical-to-basal flow) a microvessel (**Figure 8A, Figure S6A**). We observed increased HUVEC invasion, reminiscent of angiogenic sprouting [40], into the subluminal collagen I hydrogel upon application of transmural flow in the basal-to-apical direction in *PIK3CA^E542K^* microvessels as compared to control HUVECs or *PIK3CA^WT^* microvessels (**Figure 8A-C**). Consistent with previous reports [41–43], this increase in cell invasion primarily occurred with the application of basal-to-apical, but not apical-to-basal transmural flow (**Figure 8A-B**).

**Figure 8.**
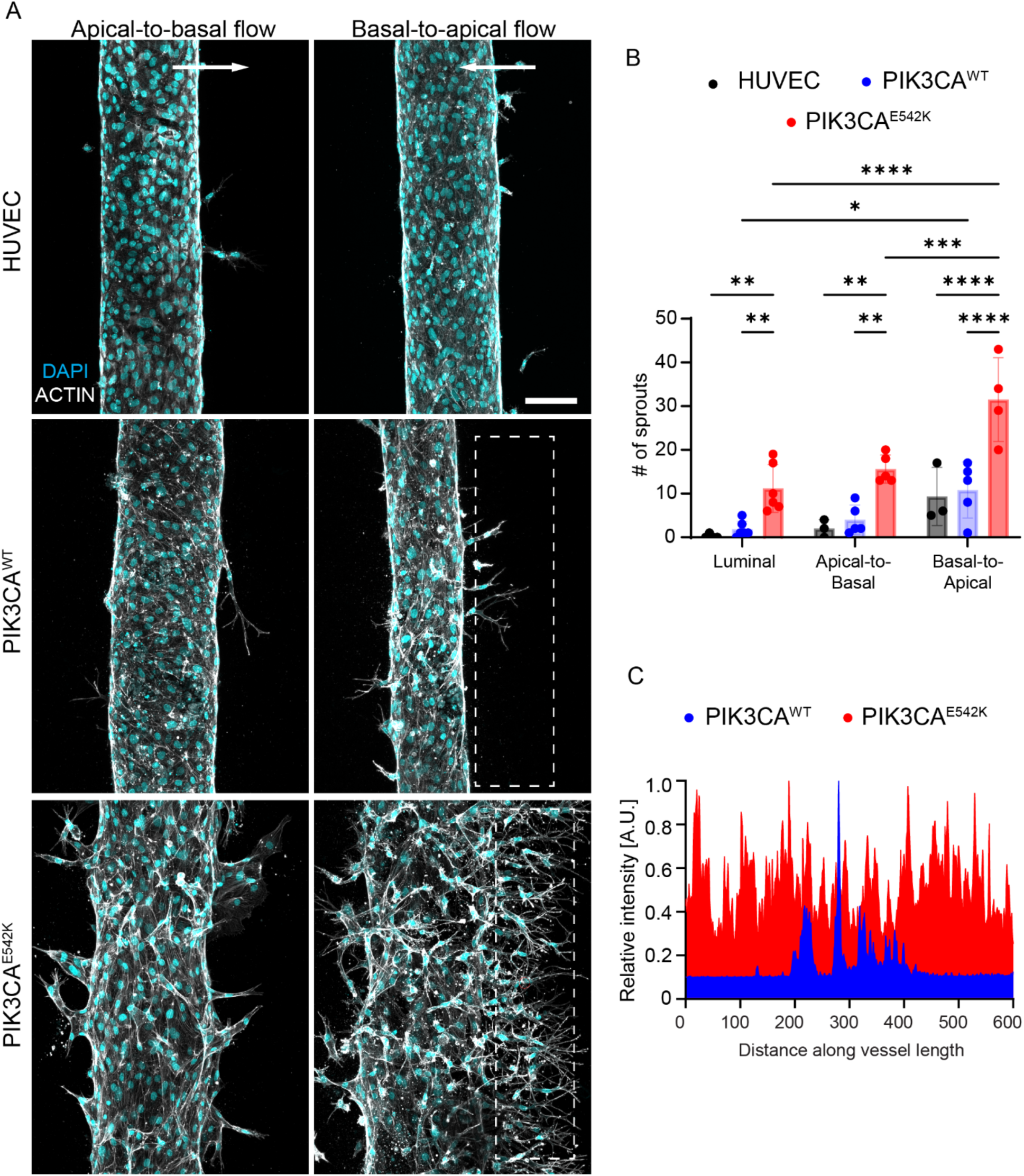
Basal-to-apical transmural flow exacerbates hypersprouting in PIK3CA mutant microvessels. (**A**) Representative images of HUVEC, *PIK3CA^WT^* and *PIK3CA^E542K^* microvessels exposed to continuous rocking for 48 hr and subsequently subjected to apical-to-basal (source) or basal-to-apical (sink) transmural flow for 24 hr. Scale bar = 100 µm. (**B**) Quantification of number of spouts in microvessels exposed to either luminal, apical-to-basal, or basal-to-apical flow (n ≥ 3; mean ± s.d., two-way ANOVA followed by Tukey test). (**C**) Fluorescence intensity profiles of actin channel were used to quantify number of sprouts in *PIK3CA^WT^* or *PIK3CA^E542K^* microvessels. Dashed white boxes indicate where intensity profiles were extracted. *p< 0.05, **p< 0.01, ***p< 0.001, ****p< 0.0001 for all plots.

### PIK3CA^E542K^ activation drives non-cell-autonomous sprouting in mosaic microvessels

Slow-flow VMs arise from sporadic somatic mutations, resulting in vascular lesions consisting of endothelial cells with and without the driving mutations. The mosaic nature of vascular lesions implies a role of non-cell autonomous signaling where mutated endothelial cells influence the behavior of neighboring non-mutated cells to amplify lesion growth and progression. To determine how *PIK3CA^E542K^* HUVECs influence non-mutated HUVECs, we generated mosaic microvessels containing HUVECs expressing histone *H2B-mCherry* and inducible *tet-ON-PIK3CA^E542K^*. We then applied basal-to-apical transmural flow and investigated whether doxycycline induced expression of *PIK3CA^E542K^* influenced sprouting and invasion behavior of control *H2B-mCherry* HUVECs (**Figure 9A**). We found that doxycycline induction increased both the number of *H2B-mCherry* and *PIK3CA^E542K^* HUVECs located within sprouts (**Figure 9B**), indicating a non-cell-autonomous effect of PI3K activation on sprouting angiogenesis. We further quantified the tip cell distribution in mosaic sprouts and observed that *tet-ON-PIK3CA^E542K^* and *H2B-mCherry* HUVECs contributed to 40% and 60% of tip cells, respectively in untreated mosaic microvessels (**Figure 9C-D**). Doxycycline treatment drastically changed the proportion of control and mutant tip cells, where *PIK3CA^E542K^* HUVECs occupied 87% of the tips and *H2B-mCherry* HUVECs contributed mostly as stalk cells in mosaic sprouts of doxycycline treated microvessels (**Figure 9C-D**). Together, these results suggest that *PIK3CA^E542K^* endothelial cells are more migratory and have a propensity to lead the sprouting and invasion of non-mutated endothelial cells.

**Figure 9.**
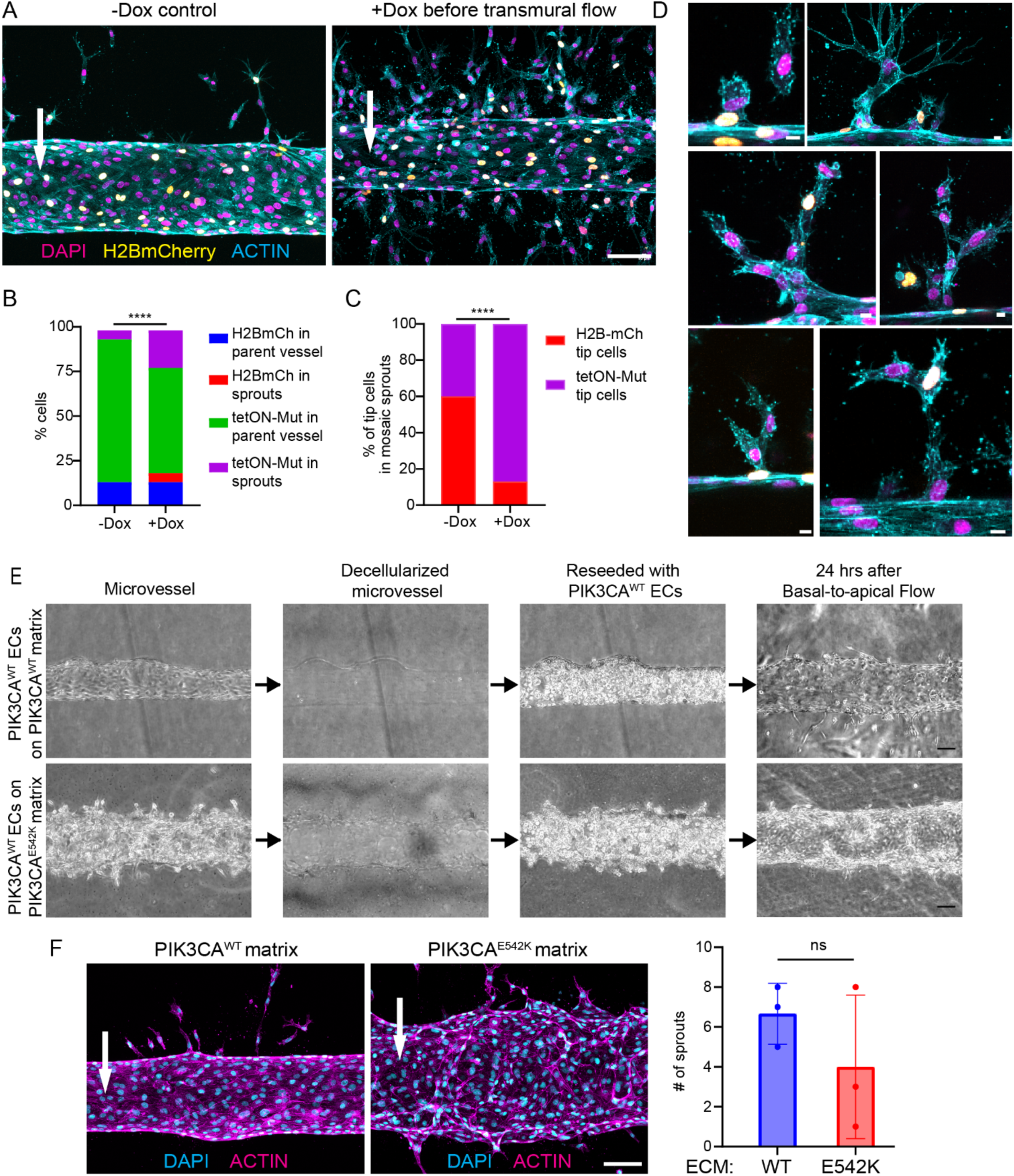
Mosaic co-culture of HUVECs with PIK3CA^E542K^ ECs resulted in increased sprouting of non-mutated HUVECs. (**A**) Representative images of mosaic microvessels generated using HUVECs expressing H2BmCherry and inducible tet-ON-PIK3CA^E542K^. Microvessels were first exposed to continuous rocking 48 hr and then were subjected to basal-to-apical transmural flow for the next 24 hrs. PIK3CA^E542K^ expression was induced upon the application of transmural flow. Scale bar = 100 µm. (**B**) Quantification of the distribution of H2BmCherry and tet-ON-PIK3CA^E542K^ cells in mosaic vessels that were subjected to basal to apical transmural flow (n ≥ 3; chi-square test). (**C**) Quantification of percent H2BmCherry and tet-ON-PIK3CA^E542K^ tip cells in mosaic sprouts (n = 5 control and 15 doxycycline treated mosaic sprouts; chi-square test). (**D**) Images of mosaic H2BmCherry (yellow) and tet-ON-PIK3CA^E542K^ sprouts stained for DAPI (magenta) and actin (cyan). Scale bar = 10 µm. (**E**) Decellularized *PIK3CA^WT^* and *PIK3CA^E542K^* microvessels were reseeded with *PIK3CA^WT^*HUVECs. Reseeded microvessels were subjected to basal-to-apical transmural flow for 24 hr. Scale bar = 100 µm. (**F**) Representative confocal images and number of sprout quantifications of transmural flow conditioned *PIK3CA^WT^* microvessels cultured on extracellular matrix modified by control or mutant cells. Scale bar = 10 µm. (n = 3; mean ± s.d., unpaired two-tailed t-test).

We next determined whether the non-autonomous effect on sprouting is a result of matrix remodeling by *PIK3CA^E542K^* HUVECs. Proteolytic degradation of the subluminal collagen matrix is known to be required for angiogenic sprout initiation and extension [44]. Previously, we described an increase in protease secretion and ECM degradation by *PIK3CA-mutant* HUVECs [19], and we thus hypothesized that ECM degradation and remodeling by *PIK3CA^E542K^* HUVECs may prime *PIK3CA^WT^*HUVECs for angiogenic sprouting. To test this hypothesis, we formed *PIK3CA^WT^* and *PIK3CA^E542K^* microvessels and lysed cells after 24 hrs, then reseeded the resulting channels with *PIK3CA^WT^*HUVECs (**Figure 9E**). We then applied basal-to-apical transmural flow and quantified the number of sprouts formed in primed ECM (**Figure 9F**). We found that *PIK3CA^WT^* HUVECs cultured in microvessels primed by *PIK3CA^E542K^* HUVECs failed to reproduce the hypersprouting phenotype (**Figure 9F**). This finding suggests that changes in matrix architecture and mechanics caused by *PIK3CA^E542K^* HUVECs are not sufficient to account for the etiopathology of VMs.

### PIK3CA activation leads to increased endothelial cell and nuclear compliance

Increased cell and nuclear compliance has been shown to promote 3D migration in various cell types [45], and we thus hypothesized that the hypersprouting observed in *PIK3CA^E542K^* microvessels was driven by changes in cell and/or nuclear mechanical properties of *PIK3CA^E542K^*HUVECs. To distinguish changes in cell intrinsic mechanics from tissue mechanics, we quantified cellular and nuclear deformation on a single cell basis using a microfluidic micropipette aspiration assay [46, 47] (**Figure 10, Figure S7**). The microfluidic device was designed to have two inlets allowing for the perfusion and loading of cells through the main channel. The main channel contains an array of 18 pockets with narrow constriction micropipette channels and are connected to an outlet port which is open to the atmosphere (**Figure S7**). A pressure of 7.0 kPa was applied to inlet 1 and 1.4 kPa to inlet 2, and both outlets are drained through constricted channels to atmospheric pressure (**Figure S7B**). Cells are initially trapped in larger pockets before deforming into 5 μm × 5 μm constricted channels under a fixed pressure gradient, and deformation was imaged via optical microscope for 2 minutes (**Figure 10B, Figure S7**). Interestingly, while untransduced HUVECs maintained cellular integrity for the duration of the experiment, *PIK3CA^WT^*and *PIK3CA^E542K^* endothelial cells were prone to fracture as the cells passed through the narrow constriction, with *PIK3CA^E542K^*cells having the lowest probability of survival throughout the experiment (**Figure 10C**). The cytoplasm and nuclear protrusion length were measured as a function of time, and while the cytoplasm of *PIK3CA^WT^* and *PIK3CA^E542K^*HUVECs deformed similarly in response to the applied pressure gradient, both cell types demonstrated greater cytoplasmic deformation and cytoplasm fracture than untransduced HUVECs (**Figure 10C-D**). The nuclei of the three cell types deformed at different rates, with untransduced HUVEC nuclei experiencing the least deformation over time and *PIK3CA^E542K^* expressing nuclei experiencing the greatest deformation (**Figure 10E**). Nuclear and cytoplasm viscoelastic properties were computed from the protrusion curves, as described previously [47]. Interestingly, *PIK3CA^E542K^* and *PIK3CA^WT^* HUVECs share similar cytoplasmic viscoelastic properties and are more compliant and less viscous than the cytoplasm of untransduced HUVECs (**Figure 10F**). The nuclei of *PIK3CA^E542K^*HUVECs showed reduced elasticity as compared to the nuclei of *PIK3CA^WT^* and untransduced HUVECs but exhibited the same viscous properties as *PIK3CA^WT^* nuclei (**Figure 10G**). Thus, *PIK3CA^E542K^* HUVECs were distinguished only by their reduced nuclear elastic modulus compared to *PIK3CA^WT^* HUVECs. Together, these results demonstrated increased cellular and nuclear compliance in PIK3CA-mutant ECs which may underlie dilation and overgrowth of venous and lymphatic vessels.

**Figure 10.**
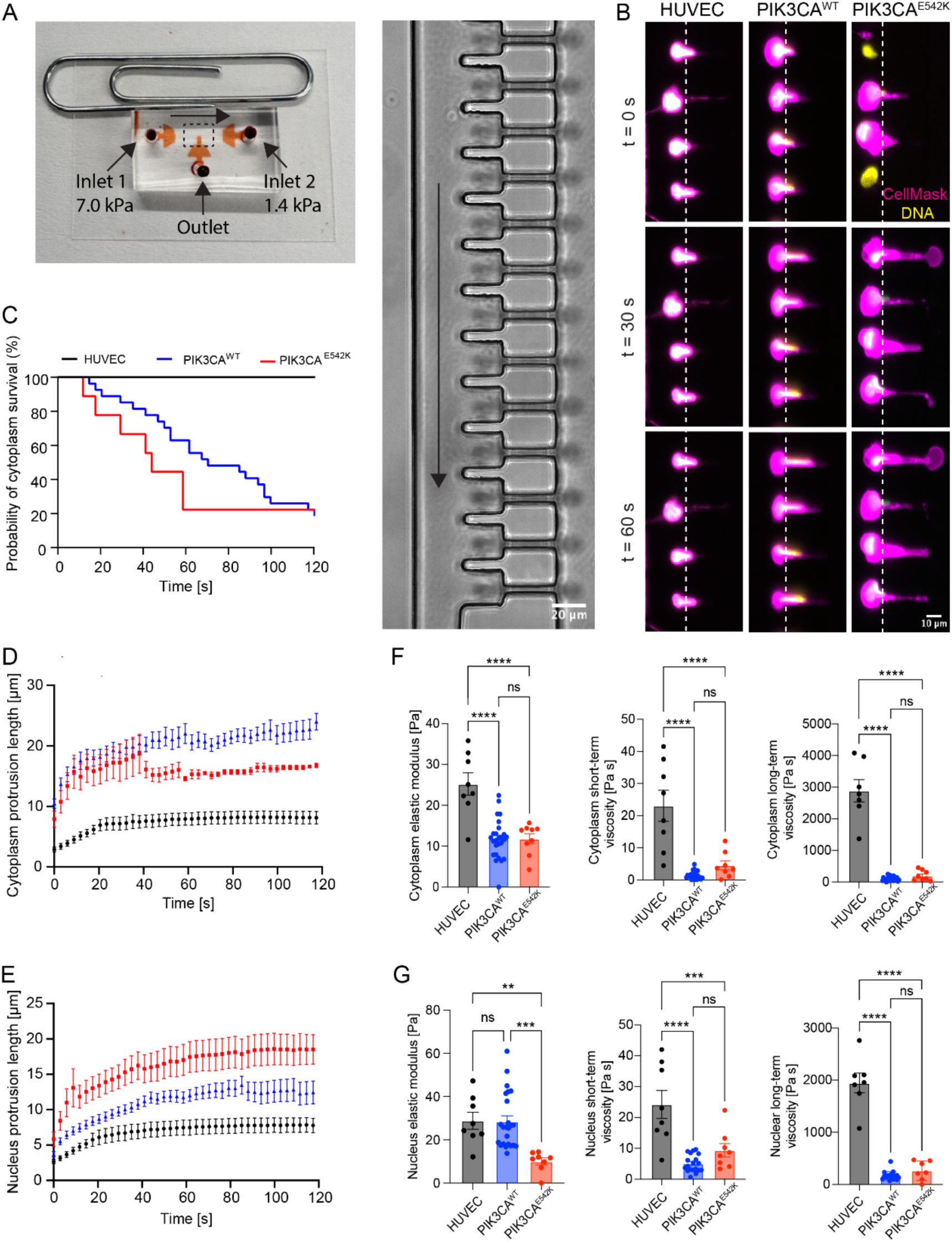
Altered cytoplasmic and nuclear mechanical properties of single cells expressing *PIK3CA^WT^*or *PIK3CA^E542K^* activating mutations. (**A**) An image of the microfluidic micropipette aspiration device labeled with an outlet and two inlets at differential pressures causing a flow vector between the two inlets. A magnified image depicts the dashed area, showing the array of pockets and constriction channels which trap and deform flowing cells under a constant applied pressure gradient. (**B**) Time-lapse images of nucleus (Hoechst) and cytoplasm (CellMask) deformation of untransduced HUVECs and endothelial cells expressing *PIK3CA^WT^* or *PIK3CA^E542K^* activating mutations. (**C**) Percentage of HUVECs that withstood deformation, as measured by the number of endothelial cells with intact cytoplasm after passage through the constriction channel relative to the total number of cells that entered the channels. (**D**) Protrusion length over time of endothelial cell cytoplasm into the constriction channels (mean ± s.e.m.). (**E**) Quantification of nuclear deformation over time of endothelial cells into constriction channels (mean ± s.e.m.). (**F**) Quantification of viscoelastic properties of the cytoplasm of endothelial cells (mean ± s.e.m., one-way ANOVA). (**G**) Elastic modulus, short-term viscosity, and long-term viscosity of the nuclei of HUVECs and endothelial cells expressing *PIK3CA^WT^* or *PIK3CA^E542K^*activating mutations (mean ± s.e.m., one-way ANOVA).

## Discussion and conclusion

*PIK3CA-*driven VMs stereotypically present in soft tissues with low rates of perfusion [4, 48]. Here, using a series of engineering and microfluidic tools, we investigated the role of the biophysical microenvironment in the development of slow-flow vascular malformations to test the hypothesis that low hemodynamic shear stress potentiates lesion formation. Alignment and elongation of the cytoskeleton in the direction of flow is a hallmark of endothelial mechanotransduction [49], and laminar shear stress has been shown to regulate vascular barrier function locally and acutely [31, 50]. Furthermore, a lack of alignment and barrier function in response to hemodynamic shear stress is indicative of endothelial dysfunction [51]. Here, using both an orbital flow system to impart a range of shear stress magnitudes and a Hele-Shaw flow cell for unidirectional, laminar flow, we observed that HUVECs and LECs expressing a constitutively active PIK3CA-activating mutation fail to elongate and align in response to laminar shear stress (**Figures 1-2**). The lack of cell alignment was also reflected in a more isotropic distribution of microtubules and intermediate filaments in mutant cells (**Figures S3-S4**). Furthermore, the permeability of PIK3CA-mutant endothelial cells remained elevated even in the presence of physiologic shear stress (**Figures 1-2**). Together, these results suggest that dysfunctional mechanotransduction and resultant insensitivity to flow could be a key mechanism that underlies the dysregulated vascular networks characteristic of VMs (**Figure S8**).

We observed that *PIK3CA^E542K^* microvessels generate lower net traction forces than *PIK3CA^WT^* microvessels (**Figure 6**). Previous work has demonstrated that in multicellular systems, traction forces measured at cell-ECM interfaces are impacted by changes in cell-cell and cell-matrix adhesion [52], and that maturation of adherens junction complexes requires a balance of MLC-mediated cell-generated forces and Rac1 activity [53]. Previously, we identified that *PIK3CA* activating mutations result in excess Rac1 activity [19]. Here, we observed that expression of *PIK3CA^E542K^*reduced MLC phosphorylation and recruitment of vinculin to adherens junction complexes, resulting in the discontinuity of junctional VE-cadherin and increased permeability (**Figure 3-4**). Together, these results suggest that pathophysiologic PI3K activation results in adherens junction instability through mechanical imbalances at cell-cell junctions, consistent with previous work demonstrating that PIK3CA signaling inhibits NUAK1-dependent phosphorylation of myosin phosphatase targeting-1 in ECs, leading to reduced overall actomyosin contractility [18]. Collectively, these results suggest that adherens junction instability and lack of maturation in response to hemodynamic signals render PIK3CA mutant cells prone to angiogenesis and could underlie the overconnected topologies characteristic of VMs.

Clinically, VMs are described as soft and compressible, and histologically they appear as a single layer of endothelial cells in enlarged vein-like channels with disorganized pericellular matrix [4]. To recapitulate the compliant ECM characteristic of tissues susceptible to VMs, we developed a 3D microfluidic model of vascular malformations and observed cellular signatures consistent with the clinical presentation of VMs, including irregular and dilated microvessels (**Figure 6**). In previous work, we performed a broad characterization of MMPs and fibrinolytic enzymes in conditioned media from wild type and mutant ECs, and we observed upregulation of MMP-1, which degrades collagen type I [19]. These results are consistent with the increased diameter of mutant microvessels (**Figure 6A-B**), and could contribute to the increased sprouting in response to flow, as perfusion has been shown previously to regulate angiogenesis in an MMP-1-dependent manner [54].

Previously, we have shown that reducing hemodynamic shear stress leads to rapid adherens junction dissolution [31], and experiments conducted in developmental animal models have demonstrated that reduction or interruption of flow induces vessel regression and network pruning [55], which is thought to be essential for optimization of efficient delivery of oxygen and nutrients [56, 57]. To recapitulate the hemodynamic signals associated with pruning, we exposed microvessels to flow for 24 hrs, then interrupted flow and cultured microvessels in static conditions for 24 hrs. Interestingly, PIK3CA mutant microvessels did not respond to flow interruption, and mutant ECs remain attached and continue to grow under static culture conditions, whereas wild-type ECs delaminate from the vessel wall (**Figure 7, Figure S5**). These results suggest that cells expressing activating PIK3CA mutations are primed for survival in the unique hemodynamic environment of slow-flow lesions, and that this survival results in a lack of pruning in response to reduced or interrupted perfusion to allow VM progression in the slow-flow microenvironment.

Computational modeling and a limited number of experiments have demonstrated that changes in vascular network topology modulate not only local wall shear stress but also luminal pressure [58]. To determine whether gradients in fluid pressure could play a role in lesion formation, we fabricated microfluidic devices to expose wild-type and mutant microvessels to transmural pressure gradients. We observed that transmural pressure gradients and resultant flow in the basal-to-apical direction results in increased sprouting and invasion of PIK3CA-mutant ECs as compared to control microvessels (**Figure 8**). Previously, immature endothelial monolayers, characterized by discontinuous junctional VE-cadherin, have been shown to sprout in response to lower basal-to-apical pressure drops than mature monolayers with continuous pericellular VE-cadherin [54], suggesting that the increased sprouting in PIK3CA-mutant ECs could be driven by immature adherens junctions. Furthermore, basal-to-apical transmural flow has been shown to synergize with VEGF gradients to increase angiogenic sprouting [42], and a recent study has demonstrated increased paracrine VEGF-C signaling in *PIK3CA*-driven mosaic LM mouse model [26]. Together, these results suggest that transmural pressure gradients and flow are a pro-angiogenic stimulus that drives lesion elaboration and growth.

Permeability measurements and images of pericellular VE-cadherin (**Figures 1, 3**), suggest that adherens junctions in mutant cells are immature and discontinuous. To relate the junctional phenotypes to the dysfunctional response to shear stress, we investigated whether downstream effector signaling modulated by key mechanosensory complexes that contain VE-cadherin [22, 23, 31] were differentially regulated in mutant cells. We first quantified the stability of VE-cadherin intercellular adherens junctions using a pulse-chase approach and observed a significantly reduced level of labeled VE-cadherin at cell-cell contacts (**Figure 4**). We then tested the hypothesis that Src kinase, which is activated by the VE-cadherin/PECAM/VEGFR2 mechanosensory complex [23], is differentially phosphorylated by mutant cells. Interestingly, we found that Src phosphorylation was similar in magnitude and temporal activation among untransduced HUVEC, *PIK3CA^WT^*, and *PIK3CA^E542K^* cells (**Figure 4)**. Furthermore, closer inspection of the pulse-chase staining of VE-cadherin reveals an increase in the intracellular puncta of AF647-VE-cadherin, suggesting increased VE-cadherin internalization and recycling. Previous work has established mature adherens junctions as sites of cytoskeletal organization [59], that VE-cadherin is necessary for proper alignment of the cytoskeleton in response to shear stress [60], and that pericellular VE-cadherin localization balances cell-generated forces associated with cortical actin elaboration [53, 61]. Thus, our data suggest that hyperactive PIK3CA activity, driven by mutant expression, reduces adherens junction stability through increased VE-cadherin turnover, which results in defective coordination of cytoskeletal dynamics and resultant alignment in response to flow, but that sites of assembled VE-cadherin are capable of transducing flow responses as measured by Src phosphorylation. Future work will seek to corroborate these results and provide greater mechanistic insight through dynamic characterization of gene expression profiles and direct testing of mechanosensory complex assembly and stability through immunoprecipitation.

Previous studies have demonstrated that nuclear stiffness and deformability play a role in 3D cellular migration through dense collagen matrices and spatially constraining micropores [45, 62]. Using a microfluidic micropipette aspiration assay, we found that *PIK3CA^WT^* and *PIK3CA^E542K^* ECs exhibit increased cellular compliance compared to untransduced HUVECs (**Figure 10**) and that *PIK3CA^E542K^* ECs have significantly more compliant nuclei than *PIK3CA^WT^* ECs. The increased cellular and nuclear compliance in PIK3CA-mutant ECs may facilitate EC migration and sprout formation, potentially contributing to hypervascularization in VM lesions. Previous work has demonstrated a role of Akt in phosphorylating Lamin A/C, resulting in degradation in keratinocytes [63] and C2C12 myocytes [64]. Furthermore, *Lmna^-/-^* MEFs demonstrate increased nuclear deformation in a microfluidic micropipette aspiration assay similar to the one employed here [47]. These results are consistent with our observations that *PIK3CA^E542K^* cells, which we have previously shown have increased p-AKT (Ser^473^) as compared to GFP transduction control cells [19], are characterized by increased nuclear deformation (**Figure 10**). While the specific mechanisms that relate PIK3CA activation to nuclear compliance remain unclear, increased invasion of colorectal cancer cells expressing PIK3CA activating through 8 μm pores and through Matrigel has been observed [65]. Conversely, knockdown of PIK3CA in glioblastoma cells reduces invasion into Matrigel [66], collectively suggesting that PIK3CA-mediated changes in nuclear mechanics could underlie a wide variety of pathogenic processes.

In summary, our findings demonstrate a role of altered mechanotransduction and cell mechanics in PIK3CA-driven vascular malformations (**Figure 8**). These findings could potentially be leveraged for genetic diagnosis and treatment of vascular anomalies. Molecular genetic testing for VMs remains challenging as the lesions are mosaic, with only a small population of cells expressing the driving mutations. Therefore, these reported changes in cellular mechanics may potentially be used as a biomarker for the sorting and enrichment of mutated ECs from patient biopsies. Further, targeting the defective contractility and recalibrating the mechanotransduction in mutant endothelial cells could represent an avenue to reduce or regress lesion progression.

## Materials and methods

### Cell culture, cloning, and lentiviral transduction

HUVECs (Lonza) and juvenile HDLECs (Promocell) were cultured in endothelial growth medium (EGM)-2 growth medium (Lonza) in a humidified incubator at 37 °C and 5% CO_2_. HUVECs were used between passages 6 and 12. HDLECs were used between passage 3 and 7. Lentivirus generation and transduction of HUVECs with pHAGE-PIK3CAWT, pHAGE-PIK3CAE542K, and pLenti6-H2B-mCherry (Addgene, plasmid #89766, gift from Torsten Wittmann) were performed as previously described [19]. PIK3CAE542K cDNA was amplified from pHAGE-PIK3CAE542K (Addgene, plasmid # 116479, gift from G. Mills and K. Scott) and was subcloned into pCW57.1, a third-generation inducible lentiviral expression vector (Addgene, plasmid # 41393, gift from David Root). Q5-site directed mutagenesis (New England Biolabs) was performed on pHAGE-PIK3CAE542K to generate pHAGE-PIK3CAWT plasmid. All plasmids were expressed through lentiviral transduction. For lentiviruses generation, HEK-293T cells were transfected with pHAGE or pCW57.1 constructs, pSPAX2 plasmid, and pMD2.g plasmid (Adggen plasmid # 12260 and #12259, gifts from D. Trono) using calcium phosphate transfection method. Lentiviral supernatants were collected between 48 hours to 72 hours post transfection. Supernatants containing lentiviruses were filtered purified using 0.45 µm PES filters. Cytoplasmic GFP fluorescence signal through the expression of pHAGE-PIK3CAE542K-ires-GFP or pHAGE-PIK3CAWT-ires-GFP was used to measure transduction efficiency and lentivirus tittering assays were performed to determine the lowest concentration required to achieve >90% transduction efficiency in HUVECs. HUVECs expressing pCW57.1-tetON-PIK3CAE542K were selected with 2 µg/mL of puromycin (Corning) and 5 µg/mL of blasticidin (Fisher) in EGM-2, respectively for at least two cell passages. Doxycycline (Sigma, 10 µg/mL in EGM-2) was used to induce the expression of tet-ON-PIK3CA^WT^ or tet-ON-PIK3CA^E542K^.

### Hemodynamic shear stress via orbital shaking and 2D permeability assay

HUVECs were seeded at 0.15 x 10^6^ cells per well in a 6-well plate (Corning Costar 6-well Clear TC-treated, Cat #3516) coated with 50 µg/mL of rat-tail derived type I collagen (Corning 354326) in 0.02 M acetic acid solution. Cells were allowed to adhere overnight prior to the application of orbital shaking. Fresh EGM-2 (2 mL) was added into each well prior to the application of orbital flow. Orbital flow was applied by culturing ECs for 3 days on an orbital shaker set to 200 rpm, which corresponds to a maximum wall shear stress of 15 Dyn/cm^2^ [67]. Computational fluid dynamics (CFD) models verified with particle imaging velocimetry (PIV) demonstrate that a well of this geometry, fill volume, and rotational speed will generate periodic wall shear stresses with mean values spanning from 0.8 – 13 Dyn/cm^2^ from near the center of the well to the outer edge, respectively [67]. For measurements of local changes in vascular permeability, 0.3 x 10^6^ cells HUVECs were seeded in each well of a 6-well plate coated with 50 µg/mL of biotinylated-human fibronectin (Corning 354008) in PBS. Fibronectin biotinylation was performed with EZ-link-Sulfo-NHS-LC-Biotin (Thermo Fisher A39257) as described previously [68]. Briefly, 0.1 mg/mL of fibronectin in PBS was incubated with 0.5 mM of EZ-link-Sulfo-NHS-LC-Biotin for 30 min at room temperature. At the end of orbital flow culture, cells were incubated with 25 mg/mL Cy3-Streptavidin (Jackson Immuno Research Labs 016160084) in PBS containing calcium and magnesium (PBS++) for 2 minutes. After streptavidin labeling of exposed matrix, cells were rinsed once with PBS++ and were fixed with 4% paraformaldehyde (Electron Microscopy Sciences) in PBS++ for 15 min at room temperature. Permeability was measured by quantifying the streptavidin positive area using the default auto thresholding method in ImageJ.

### Hydrogel formation and cell seeding in microvessel microfluidic devices

Microfluidic devices were fabricated as described in the *Supplementary Methods*. To improve hydrogel adhesion to PDMS devices prepared by photolithography, 2 mg/mL dopamine hydrochloride in 10 mM of pH 8.5 Tris HCl buffer (BioWorld) was pipetted into the hydrogel chamber, incubated for 1 hr at room temperature, rinsed twice with DI-H_2_O, left dry at room temperature, and sterilized via UV for 20 min. Sterile 160 µm stainless steel acupuncture needles (Llahsa Oms) were coated with 0.01% bovine serum albumin (BSA) in PBS for at least 30 min prior to hydrogel injection. BSA-coated needles were inserted into each device, and 2.5 mg/mL of rat tail-derived collagen I hydrogel was prepared and pipetted into the hydrogel chamber as described previously [37], and allowed to polymerize at 37 °C for 30 min prior to hydrating with EGM-2. After 4 hrs of further incubation at 37 °C, acupuncture needles were gently removed from devices to form a cylindrical channel within the collagen I hydrogel. Needle inlets were sealed with vacuum grease (Millipore Sigma) and fresh EGM-2 was added to the media ports of the devices. Devices were kept on a rocker at 5 cycles/min at 37 °C and 5% CO_2_ for at least 12 hrs prior to cell seeding to rinse and equilibrate the collagen I hydrogel [37]. To form microvessels, HUVECs were resuspended at 0.3 x 10^6^ cells/mL in EGM-2. Media was removed from both media ports and gel filling pipette tips were used to aspirate and empty the hydrogel ports prior to seeding. To promote pressure driven flow into the microchannel, 50 µL and 40 µL of cell suspension were added to the inlet and outlet media ports, respectively. Devices were inverted every minute for 8 min at room temperature, followed by incubating upside-down in a tissue culture incubator for 6 minutes. Devices were subsequently inverted and incubated for an additional 6 minutes in the incubator. After 20 min of seeding, devices were placed on a rocker for at least 2 hours before changing EGM-2 media, which was subsequently changed daily.

### Transmural flow in two-channel devices

Two-channel devices were used to apply transmural flow into or out of microvessels (**Figure S6**). For the application of transmural flow, microvessels were first cultured on rocker (5 cycles per minute at ± 30° angle) for 2 days. Media reservoirs were inserted into all four media ports and any gaps between reservoirs and PDMS were sealed with vacuum grease. Pre-warmed EGM-2 was added either to the empty or microvessel channel to apply a 20 mm H_2_O hydrostatic pressure driven transmural flow into or out of the microvessels, respectively. Microvessels were fixed 24 hours after the application of transmural flow.

### Microfluidic micropipette aspiration

Microfluidic devices were fabricated as described in the *Supplementary Methods.* The micropipette aspiration device was used to deform cells under an applied pressure gradient. Untransduced HUVECs, *PIK3CA^WT^*, and *PIK3CA^E542K^* cells were cultured as described previously before being stained with Hoechst 3342 (10 mg/mL, Thermo Scientific) and CellMask Deep Red plasma membrane stain (5 mg/mL, 1:2000 v/v, Invitrogen) for 10 min to stain the nucleus and plasma membrane respectively. Stained cells were washed with PBS and allowed to recover in EGM-2 for 30 minutes. Cells were then trypsinized, counted, and resuspended at a concentration of 5,000,000 cells/mL in BSA (20 mg/mL in PBS) before being put on ice. Constant-pressure microfluidic flow controllers (“Flow EZ,” Fluigent) were used to apply flow to the opposing inlets at constant pressures, creating flow from one inlet to the other using a known pressure gradient. Two flow controllers were each connected to a dry air source and a power outlet. One controller was used to drive stained and suspended cells from a 15 mL conical tube into one inlet of the microfluidic device at a pressure of 7.0 kPa. A second controller was used to drive cell-free PBS from a 15 mL conical tube into the second inlet of the microfluidic device at a pressure of 1.4 kPa. The outlet port of the device was open to the atmosphere (**Figure S7**).

### Decellularization and reseeding of decellularized microvessels

Fresh sterile decellularization solution (0.5% Triton with 50 mM NH_4_OH in PBS) was prepared on the day of decellularization. Microvessels were rinsed once with sterile PBS prior to decellularization. After PBS wash, 100 µL of decellularization solution was added to the media ports of each device. Devices were kept on a rocker at 37 °C and 5% CO_2_ for 15 min. Decellularized devices were washed three times with PBS and residual DNA was digested via treatment with 10 µg/mL of DNAseI (Sigma Aldrich, D4527) in PBS++ for 45 minutes at 37 °C. DNAseI treated devices were washed three times with PBS and were incubated overnight on a rocker in fresh EGM-2 media. Devices were reseeded with *PIK3CA^WT^* cells the next day as described in the “Hydrogel formation and cell seeding” section. Basal-to-apical pressure driven transmural flow was applied a day after reseeding as described above.

### Immunostaining

Mouse Anti–VE-cadherin (F-8) was from Santa Cruz Biotechnology. Rabbit Anti-VE-cadherin (ab33168) was from Abcam. Mouse Anti-Vinculin (hVIN-1) was from Sigma Aldrich. 4′,6-Diamidino-2-phenylindole (DAPI), rhodamine phalloidin (1 mg/ml), Alexa Fluor 488 phalloidin (1 mg/ml), and Alexa Fluor–conjugated secondary antibodies were from Life Technologies. Unless otherwise mentioned, for endpoint immunostaining, cells or microvessels were fixed in 4% PFA in PBS++ for 20 minute at room-temperature. For microtubule staining, cells were fixed in 100% Methanol for 3 minutes at -20 °C. Fixed samples were subsequently washed and rehydrated with PBS. For visualization of vinculin at focal adhesion, cells were subjected to mild permeabilization using 0.1% Triton X-100 in 4% PFA. Cells were partially perm-fixed for 5 minutes prior to switching to complete fixation in 4% PFA for 15 minutes at room-temperature. All samples were permeabilized with 0.3% Triton X-100 (Millipore Sigma) for 30 minutes at room temperature and non-specific antibody binding was blocked by treatment with 2% (w/v) BSA in PBS for an hour at room temperature. Primary antibodies against VE-cadherin (1:200, v/v, F-8, Santa Cruz Biotechnology; 1:250, v/v, ab33168, Abcam), vinculin (1: 400, v/v, hVIN-1, Sigma Aldrich), α-tubulin (1:1000, v/v, DM1A, Cell Signaling), and vimentin (1:200, v/v, D21H3, Cell Signaling) were diluted in 2% BSA blocking solution and applied overnight on a laboratory rocker at 4 °C. Cells and devices were subsequently rinsed three times over 15 min with PBS, and were incubated in secondary antibodies (1:1000, v/v, goat anti-mouse or goat-anti rabbit immunoglobulin G conjugated to Cy3 or Alexa Fluor 647, Thermo Fisher Scientific), Alexa Fluor 488 or rhodamine phalloidin (1:200, v/v, Thermo Fisher Scientific), and DAPI (1:1000, v/v, Thermo Fisher Scientific) diluted in blocking solution for an hour at room temperature. Labeled cells and devices were rinsed three times over 15 min with PBS and were stored in 0.01% Sodium Azide solution diluted in PBS until imaging. Orientation of actin fibers was quantitatively determined from images of phalloidin using the FibrilTool plugin for ImageJ [69].

### Western blots

Cell lysates were collected from cells cultured in complete media for 24 hours. Briefly, cells were washed once with ice cold PBS and lysed on ice with radioimmunoprecipitation assay buffer (RIPA buffer, Thermo Fisher Scientific) containing 1X HALT protease and phosphatase inhibitor (Thermo Fisher Scientific). Lysates were homogenized by pulse-vortexing and clarified by centrifugation at 14,000 x g for 5 min at 4 °C. The concentration of clarified lysates was quantified using Pierce BCA Protein Assay (Thermo Fisher Scientific). Lysates were then reduced with NuPAGE LDS reducing agent and dithiothreitol and denatured for 5 min at 100 °C. Denatured proteins were resolved in NuPAGE 4 to 12% bis-tris gradient gels (Thermo Fisher Scientific) and transferred to iBlot2 PVDF ministack membranes. Membranes were blocked with SuperBlock Blocking Buffer (Thermo Fisher Scientific) for an hour with rocking at room temperature and were incubated with the following primary antibodies for overnight with rocking at 4 °C: anti-phosphoSRC(Tyr416) (1:1000, Cell Signaling Technology, #2101), anti-p110α (1:1000, Cell Signaling Technology, #9272), anti-vinculin (1:10,000, Sigma-Aldrich, V9264), anti-VE-cadherin (1:1000, Santa Cruz, SC9989), anti-Beta-catenin (1:1000, Cell Signaling Technology, #8480), anti-glyceraldehyde-3-phosphate dehydrogenase (1:5000, Cell Signaling Technology, #2118), and anti-phospho-myosin light chain 2 (Thr18/Ser19) (1:1000, Cell Signaling Technology, #3674). After overnight incubation with primary antibodies, membranes were washed three times with TBS-T and were subsequently incubated with horseradish peroxidase conjugated antibodies (1:10,000, v/v, diluted in 5% milk (w/v)) for 1 hour with rocking at room temperature. Membranes were then washed three times over a period of 15 min with TBS-T and were developed via chemiluminescence using either SuperSignal West Femto Maximum Sensitivity Substrate reagent (Thermo Fisher Scientific) or Clarity Western ECL reagent (BioRad). Images of western blots were quantified using ImageJ.

### VE-cadherin pulse-chase staining

For pulse-chase experiment, HUVECs were seeded and subjected to orbital shaking as described in the Methods section. After conditioning with orbital flow for 3 days, cells were incubated with AF647-anti-VE-cadherin antibody (1:200, v/v, diluted in pre-warmed EGM-2 media, BD Biosciences, #561567) for 30 minutes orbital shaking. Unbound antibody was removed with two washes with pre-warmed EGM-2 media, and cells were cultured for additional 2 hours under orbital shaking prior to fixing.

### Imaging for Microvessels and flow-conditioned cells

Confocal z-stacks of microvessels (4 µm step size, captured from top to bottom of the microvessels) were acquired with an Olympus FV3000 laser scanning confocal microscope with a 10x U Plan S-Apo NA 0.4 air objective. Confocal z-stack images of cells (0.8 µm step size, captured from top to bottom of the samples) stained for VE-cadherin, vinculin, and cytoskeletal proteins were captured with a 30X U Plan S-Apo NA 1.05 silicon oil objective. Fluorescence images of cells after orbital flow were captured at 10x magnification (10X U Plan Fluor NA 0.3 air objective, Olympus) using an automated scanning stage on an Olympus wide-field microscope. Briefly, the edge of a well of a 6-well plate was positioned over the objective, and a uniform rectangular area from the edge to the middle of the well was selected. A focus map was generated to include at least 10 focus points along the length of the rectangular area. The autofocus feature was used to cycle through the focus map and set the z-focus for each imaging point used throughout the area scan. Computational image processing is described in the *Supplementary Methods*.

### Imaging for Microfluidic Micropipette Aspiration Assay

Cell deformation into the micropipette channels was imaged using a timelapse acquisition on an inverted widefield fluorescence microscope (Olympus IX83) with a white light source 20X/NA 0.45 objective. DAPI and Cy5 filters were used to image the nucleus and cytoplasm respectively, and images were acquired every 2.935 seconds for a total acquisition time of 2.5 min. At the end of each timelapse acquisition, deformed cells were cleared from the cell pockets by applying pressure at the outlet port by pipette. Cleared pockets were then filled again by flowing cells during the next acquisition period.

### Statistical analysis

Graphs were generated and statistical analyses were performed in Prism 10 (GraphPad). Unless otherwise mentioned, data were plotted as mean ± standard deviation of ≥3 experiments and data points denote values from each cell or microvessel. Data set with normal distribution were compared with two-tailed Student’s test and one-way ANOVA followed by Tukey’s post hoc test. Kruskal-Wallis test followed by Dunn’s multiple-comparison was used for comparison of non-Gaussian distributed data. Two-way ANOVA followed by Šidák’s test was used for comparison of more than two variables.

## Supporting information

Supplementary Results

## Acknowledgements

This work was funded by the National Institute of Health (R35GM142944), by the American Heart Association (CDA857738), and by research grants from the CLOVES Syndrome Community and the Lymphatic Malformation Institute. W.Y.A. is supported by grant from the CLOVES Syndrome Community. C.P.W. and E.L.D. are supported by a training fellowship from the Integrative Vascular Biology Training Program (T32HL69768), C.P.W. is supported by an American Heart Association predoctoral fellowship (24PRE1192185), and E.L.D. is supported by a Ruth L. Kirchstein predoctoral individual fellowship (F31HL162462). E.M. acknowledges support from an NIH training fellowship (T32GM135128). We thank Bob Geil for advice and help with microfabrication. The fabrication of microfluidic devices was performed in the Chapel Hill Analytical and Nanofabrication Laboratory (CHANL), a member of the North Carolina Research Triangle Nanotechnology Network (RTNN), which is supported by the National Science Foundation (ECCS-2025064), as part of the National Nanotechnology Coordinated Infrastructure (NNCI).

## Author Declarations

### Ethics Approval

Ethics approval is not required.

### Data Availability

The data that support the findings of this study are available from the corresponding author upon reasonable request.

