## Supplementary Results for "Dysfunctional mechanotransduction regulates the progression of PIK3CA-driven vascular malformations"

### Supplementary Methods

*Microvessel microfluidic device fabrication*

Silicon master molds for microfluidic devices were fabricated using photolithography as described previously [1, 2]. The same fabrication process was used for both single-channel devices and double-channel devices for transmural flow, using different film transparency masks. Briefly, after stripping the oxide layer from a 100 mm silicon wafer using dilute hydrofluoric acid (buffered oxide etchant, Transene), an adhesion layer of SU-8 2002 (Kayaku Advanced Materials) was spread using a spin coater (Laurell Technologies) at 2,000 RPM and flood exposed with UV light (SUSS Microtech) with an exposure energy of 100 mJ/cm$^2$. Subsequently, a 100 µm thick needle buffer layer was created by spinning SU-8 2050 at 1,000 RPM, and soft baking for 5 min at 65 ºC and 3.5 hrs at 95 ºC. A blocking layer consisting of 70% v/v SU-8 2010 and 30% v/v S1813 (MicroChem) was then spun onto the wafer at 1,000 RPM, and the wafer was soft baked for 30 min at 95 ºC. The needle buffer film transparency mask and wafer were loaded into a mask aligner (SUSS Microtech) and exposed to UV light at a dose of 600 mJ/cm$^2$ before a post-exposure bake for 5 min at 65 ºC and 12 min at 95 ºC. A 200 µm needle guide layer was then spun on the wafer using SU-8 2150 at 1,800 RPM then soft baked for 5 min at 65 ºC and 80 min at 95 ºC. The wafer and needle guide film transparency masks were loaded into the mask aligner and aligned with the needle buffer layer before exposing with UV light at a dose of 300 mJ/cm$^2$ before a post-exposure bake for 5 min at 65 ºC and 12 min at 95 ºC. The gel top layer was spun on the wafer using SU-8 2150 at 1,400 RPM then soft baked for 5 min at 65 ºC and 80 min at 95 ºC. The gel top transparency mask and wafer were loaded into a mask aligner, aligned with the needle guide layer, and exposed to UV light at a dose of 300 mJ/cm$^2$ before a post exposure bake for 5 min at 65 ºC and 12 min at 95 ºC. The wafer was allowed to cool to room temperature and was developed for 10 minutes in SU-8 developer (Kayaku Advanced Chemicals).

*Microfluidic micropipette aspiration device*

Microfluidic devices were designed and fabricated as previously described [3]. The critical dimensions of the device involve two fluid inlets converging onto 18x cell pockets that were 20 µm long, 15 µm wide, and 15 µm high (**Figure S7**). As cells flowed into the device, individual cells were trapped in each pocket and were deformed under a pressure gradient into micropipette channels designed to be 5 µm on all sides (see Microfluidic micropipette aspiration section below) (**Figure S7**). Silicon wafers (76.2 mm, University Wafer) were placed into dilute HF for 1 min to strip the oxide layer. To create the micropipette channels, wafers were spin-coated with SU-8 2005 (Kayaku Advanced Chemicals) at 3,000 RPM to a first-layer thickness of 5 µm. Wafers were soft baked for 2 minutes at 95 ºC. Using a mask aligner (SUSS Microtec), wafers were exposed to UV light (112 mJ/cm$^2$) through a 365 nm UV filter and a chrome mask of the micropipette channels. Wafers were post-exposure baked for 3 min at 95 ºC and were cooled to room temperature before being developed in SU-8 Developer (Kayaku Advanced Chemicals) for 1 minute on an orbital shaker at 100 RPM. Wafers were rinsed in isopropanol. To create the cell pockets and fluid inlets, a layer of SU-8 2007 was spun onto the developed wafers at 3,000 RPM to a second-layer height of 15 µm. Wafers were soft baked for 2.5 minutes at 95 ºC. Using a mask aligner, wafers were exposed to UV light (184 mJ/cm$^2$) through a 365 nm UV filter and a chrome mask of the fluid inlets and cell pockets. Wafers were post-exposure baked for 3.5 min at 95 ºC and were cooled to room temperature before being developed in SU-8 Developer (Kayaku Advanced Chemicals) for 2.5 minutes on an orbital shaker at 100 RPM.

*Soft lithography*

Silicon wafers were prepared for soft lithography through plasma treatment for 30 seconds followed by passivation by vapor deposition of Trichloro(1*H*, 1*H*, 2*H*, 2*H*-perfluorooctyl)silane (Millipore Sigma) overnight. Individual devices were molded from passivated silicon wafers using standard soft lithography techniques. Polydimethyl siloxane (PDMS, Sylgard 184, Dow Corning) was mixed at a 10:1 ratio of base:crosslinker and was degassed for 1 hr in a vacuum chamber before being poured onto the wafers. Wafers with uncured PDMS were degassed for 30 min before being cured overnight at 65 ºC. The first 3 PDMS molds of devices were discarded due to residual silane on the PDMS surface. For use in bonded devices, PDMS was cut into individual devices using a razor blade, and holes for the fluid inlets and outlets were punched biopsy punches. PDMS devices and glass slides were cleaned using clear tape and isopropanol. The devices and glass slides were treated with oxygen plasma (Harrick Plasma) for 2.5 min before bonding. For the micropipette aspiration system, bonded devices were annealed for 7 minutes at 100 ºC and passivated with sterile bovine serum albumin (BSA, 20 mg/mL in PBS).

*Quantification of cell alignment*

A custom Cell Profiler workflow combining Stardist and Cellpose deep-learning based image segmentation methods was used to quantify cell shape and cell alignment. Briefly, a Stardist model was used on DAPI channel for nuclear segmentation and a pre-trained Cellpose model was used to on DAPI and VE-cadherin channels to identify outlines of individual cells. The shape and orientation of segmented cells were measured using the MeasuredObjectSizeShape function in Cell Profiler.

*Quantification of actin, microtubule, and vimentin cytoskeleton orientation*

Segmented cellular outlines and maximum intensity projections of confocal z-stack images were used for fibril analysis. Cytoskeletal fibers orientation was calculated using the FibrilTool Batch plugin in ImageJ [4]. Briefly, binary images of individual cells were exported from Cell Profiler, and regions of interest defining the outline of each segmented cells were extracted using the "Analyze Particles" and "Add to Manager" functions in ImageJ. Roi sets and maximum intensity projection images were subsequently loaded into the FibrilTool macro for analysis of fibrils orientation within each cell.

*Quantification of percent cellular area occupied by actin, microtubule, and vimentin cytoskeleton*

Sum intensity projections of confocal z-stack images of actin, microtubule, or vimentin cytoskeleton were used for image analysis. Image thresholding and analysis were performed in CellProfiler. Adaptive two-class Otsu thresholding algorithm using a 20 pixels (0.414 micron/pixel) adaptive window, without smoothing were used for thresholding.

*Quantification of internalized pulse-labeled VE-cadherin puncta*

Maximum intensity projections of confocal z-stack images of pulse-labeled VE-cadherin were used for image analysis. Confocal z-stacks of microvessels (0.8 μm step size, captured from top to bottom of the microvessels) were acquired with an Olympus FV3000 laser scanning confocal microscope with a 30x U Plan S-Apo NA 1.05 silicon oil objective. Image thresholding and analysis were performed in CellProfiler. Manual thresholding with the same value were used across all samples.

*Quantification of Vinculin overlap and fluorescence intensity at cell-cell border*

The segmented binary cytoplasm was first shrunk by 5 pixels. The difference between the original cytoplasm and the shrunken cytoplasm objects was compared and identified as cell-cell boundary using the IdentifyTertiaryObjects function in Cell Profiler. Vinculin within cell-cell boundaries were segmented using manual thresholding set at 0.45 and the area occupied by segmented vinculin at each cell-cell boundary were quantified. Percent vinculin overlap was defined as area of segmented vinculin at cell-cell boundaries relative to area of cell-cell boundary of each cell. Mean fluorescence intensities of vinculin and VE-cadherin at cell-cell boundaries were measured using the MeasureObjectIntensity function in Cell Profiler.

*Vessel sprout and diameter quantification*

Confocal z-stacks of DAPI and actin channels (4-μm step size) were captured from bottom to top of the vessel-on-chip. Maximum intensity z-projection of actin channel was generated, and a rectangular area aligned with the top edge of the vessel was selected using the rotated rectangle tool in ImageJ. An intensity profile of maximum-projected actin channel was generated using the plot profile function in ImageJ. Peaks with a gray value intensity greater than 200 were counted as vessel sprouts, and the process was repeated for the bottom edge of the vessel. The sprout count was manually verified. For quantification of vessel diameter, a bounding rectangle tool was used to manually fit the horizontal length of the vessel and the length of the bounding box was extracted as vessel diameter.

*Quantification of cell and nuclear deformation*

Nuclear and cytoplasm deformation of cells in the micropipette aspiration device were measured in ImageJ. A stack containing only the micropipette channels was isolated using a rectangular crop. Each cell was individually cropped, and cells that did not fully plug the pockets were excluded from analysis. Videos of each cell being deformed through the micropipette were analyzed by splitting the color channels to isolate the nuclear and cytoplasm stain respectively. Then, a threshold was used to create binary videos of the deforming nucleus or cytoplasm against a black background. Lastly, the “Analyze Particles” feature was used to measure each cell as a box with an increasing width as the nucleus or cytoplasm deformed down the length of the micropipette channel. Protrusion length was defined as the distance the cytoplasm or nucleus displaced into the constricted channel, and protrusion length as a function of time was fit for each individual cell and nucleus was fit to a Jeffreys model of a spring and dashpot in series [3] using MATLAB according to Eqs. 1-2:

$$L(t) = \frac{R_{eff}\Delta P}{E}\left(1 - e^{\frac{-E}{3\pi\eta_1}t}\right) + \frac{R_{eff}\Delta P}{3\pi\eta_2}t \qquad [1]$$

$$R_{eff}^4 = \frac{2}{\pi}\frac{W\times H^3}{\left(1+\frac{H}{W}\right)^2 + f'}, \qquad [2]$$

Where $L(t)$ represents the measured protrusion of cytoplasm or nucleus, $\Delta P$ represents the applied pressure drop by the constant-pressure pumps, $t$ represents time in seconds, and $W$ and $H$ represent the width and height of the constriction channel, used to calculate the effective radius $f' = 0.5928$ [5].

The first term in the Jeffrey’s model equation dominates at short time scales and plots as a rapidly rising curve, governed by the elastic modulus $E$ and short-term viscosity $\eta_1$. The second term in the Jeffrey’s model equation plots linearly and is governed by long-term viscosity

$\eta_2$ [3]. From the fit, the elastic modulus, short-term viscosity, and long-term viscosity for each cell and nucleus was determined and plotted in Prism.

### Supplementary Results

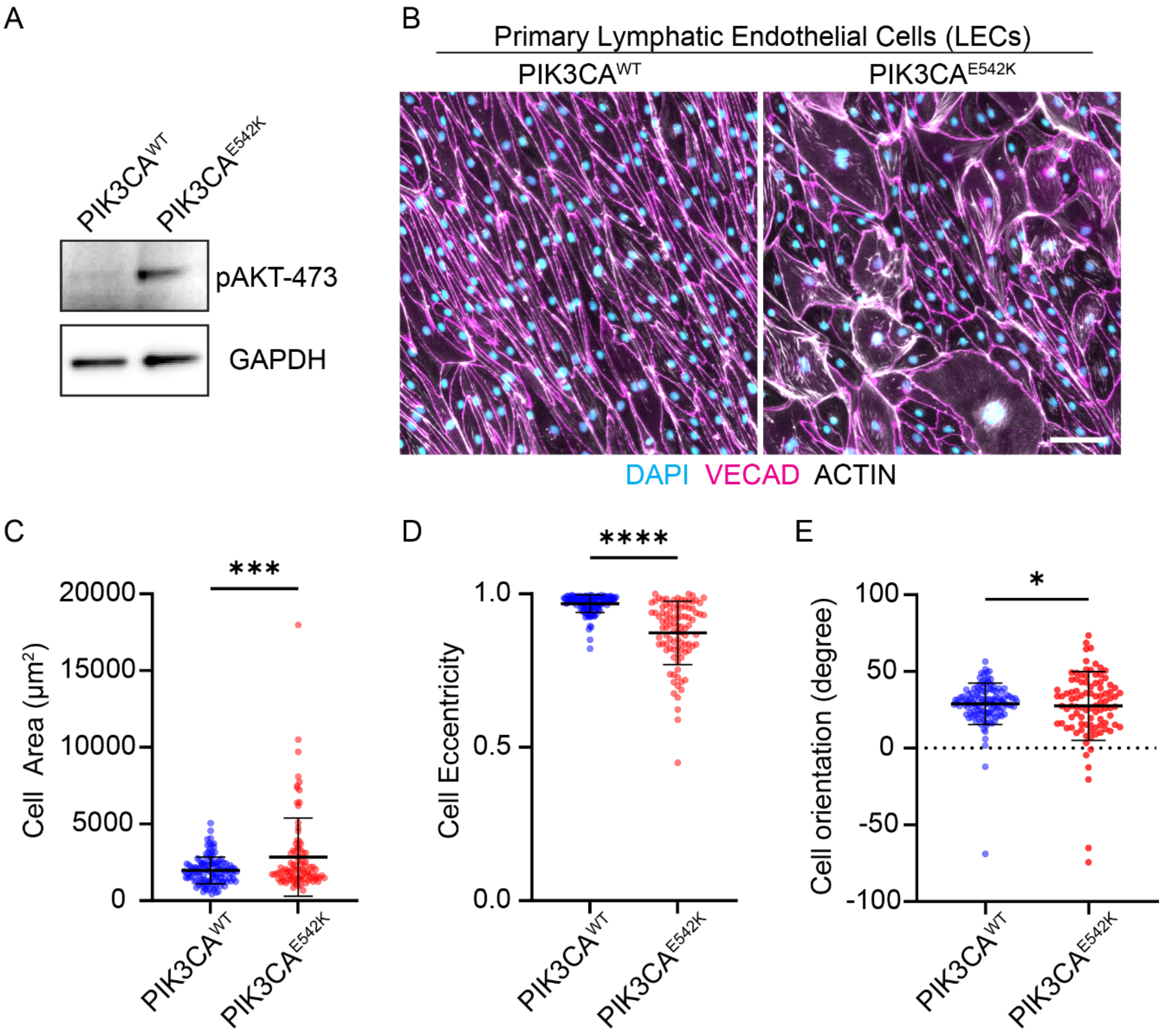


**Figure S1. Defective mechanotransduction of hemodynamic shear stress in lymphatic endothelial cells expressing *PIK3CA*$^{E542K}$.** (**A**) Phospho-AKT (Ser473) and GAPDH protein levels in lymphatic endothelial cells (LECs) expressing *PIK3CA*$^{WT}$ or *PIK3CA*$^{E542K}$. (**B**) Representative images of *PIK3CA*$^{WT}$ and *PIK3CA*$^{E542K}$ LECs cultured for 72 hrs under orbital shaking conditions. Images taken near the edge of the well, scale bar = 100 µm. (**C-E**) Quantification of cell area (**C**), cell eccentricity (**D**), and cell orientation (**E**) of flow-conditioned *PIK3CA*$^{WT}$ and *PIK3CA*$^{E542K}$ LECs.

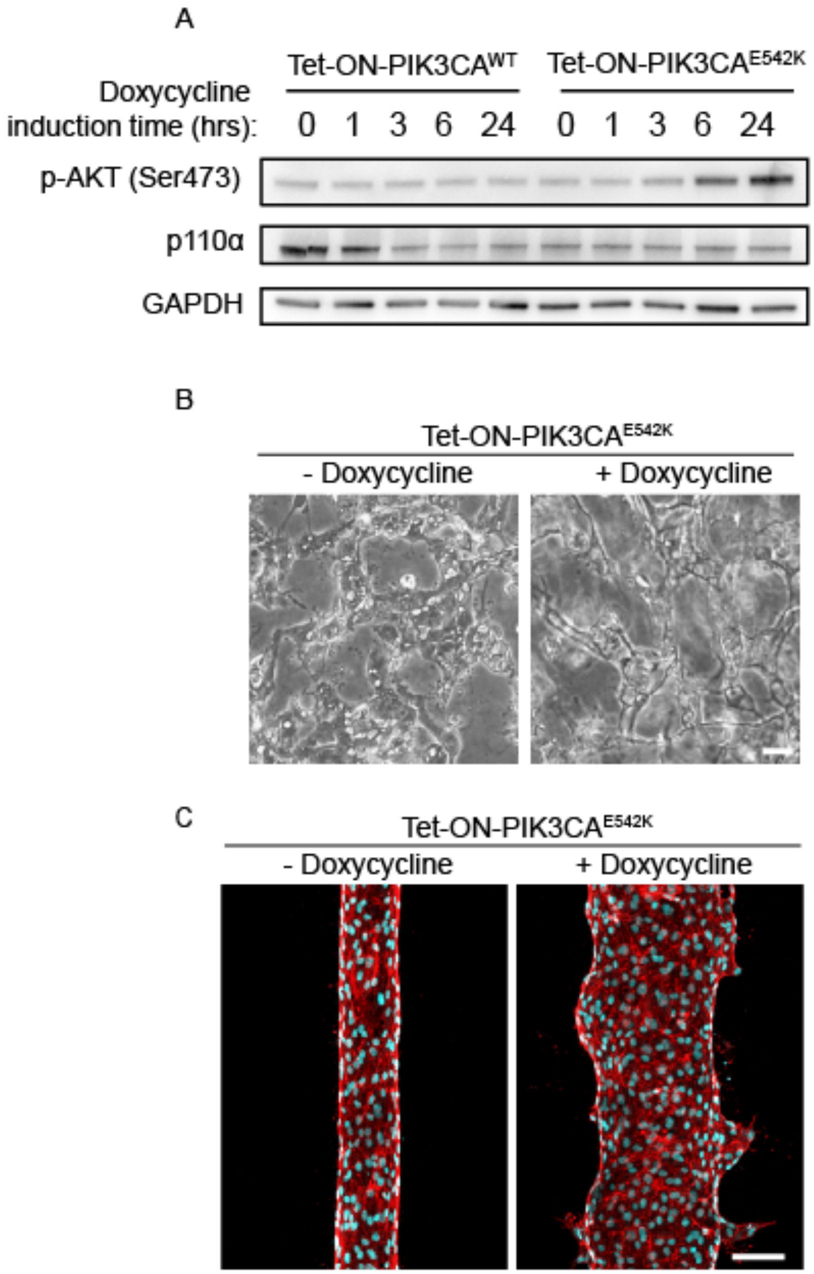


**Figure S2. Generation and characterization of *Tet-ON-PIK3CA$^{E542K}$* HUVECs.** (**A**) phospho-AKT (Ser473), p110α, and GAPDH protein levels after doxycycline-induced expression of PIK3CA$^{WT}$ and PIK3CA$^{E542K}$ in HUVECs. (**B**) Effect of doxycycline induction on *Tet-ON-PIK3CA$^{E542K}$* microvascular network formation. Scale bar = 10 μm. (**C**) Control and doxycycline treated *Tet-ON-PIK3CA$^{E542K}$* microvessels. Scale bar = 100 μm.

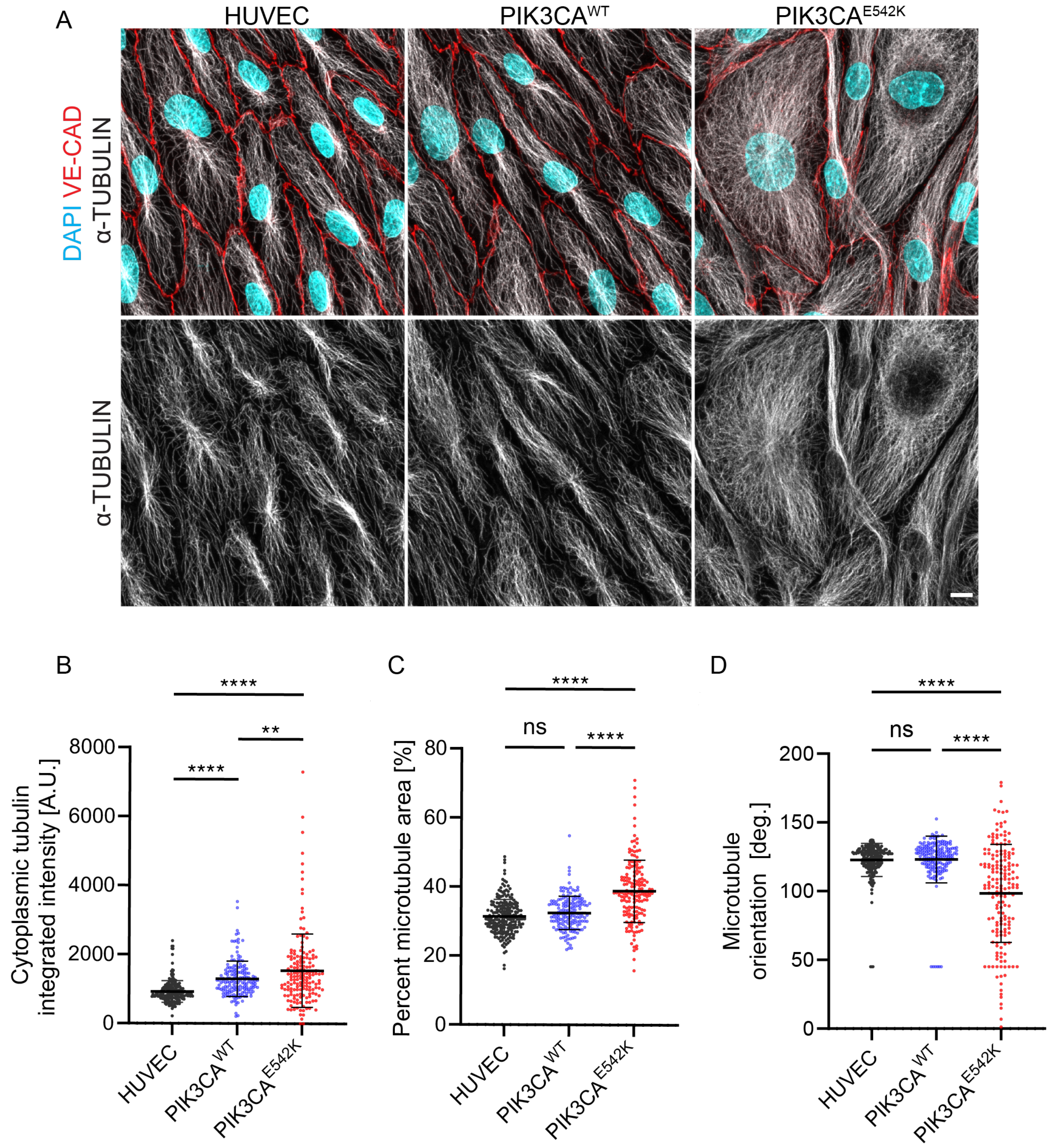


**Figure S3. Expression of *PIK3CA$^{E542K}$* increased microtubule density and impaired shear stress-mediated microtubule alignment.** (**A**) Representative images of untransduced HUVECs or HUVECs expressing *PIK3CA$^{WT}$* or *PIK3CA$^{E542K}$*. Cells were cultured for 72 hrs under orbital shaking conditions, and stained for DAPI (blue), VE-cadherin (red), and α-tubulin (grey). Images taken near the edge of the well, scale bar = 10 μm. (**B-D**) Quantification of total microtubule intensity (**B**), percent microtubule area (**C**), and microtubule orientation (**D**) within each cell.

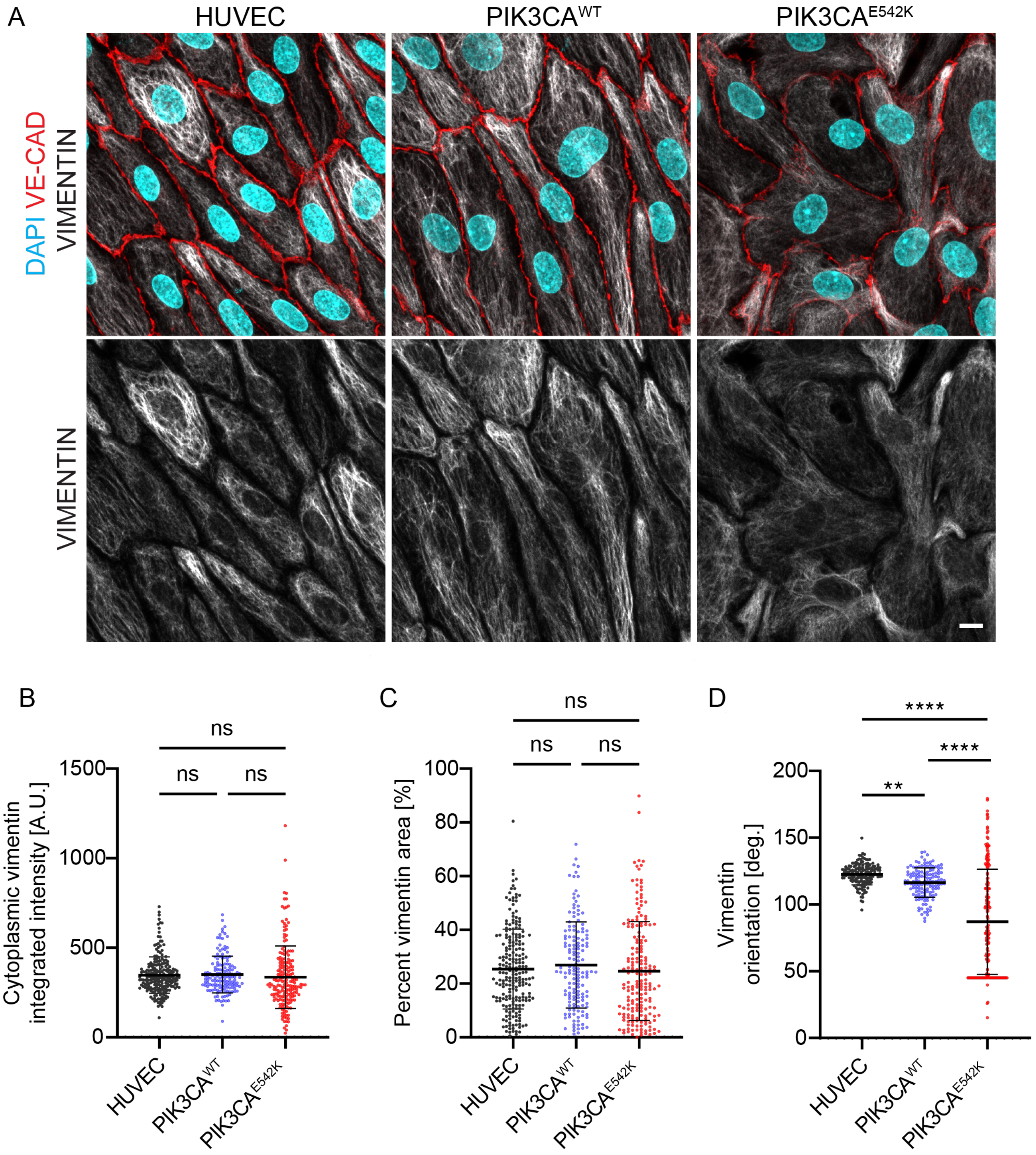


**Figure S4. Defective vimentin alignment in HUVECs expressing *PIK3CA*$^{E542K}$.** (**A**) Representative images of untransduced HUVECs, *PIK3CA*$^{WT}$ or *PIK3CA*$^{E542K}$ HUVECs cultured under orbital flow, and stained for DAPI (blue), VE-cadherin (red), and vimentin (grey). Images taken near the edge of the well, scale bar = 10 µm. (**B-D**) Quantification of total cytoplasmic vimentin intensity (**B**), percent vimentin area (**C**), and vimentin orientation (**D**) within each cell.

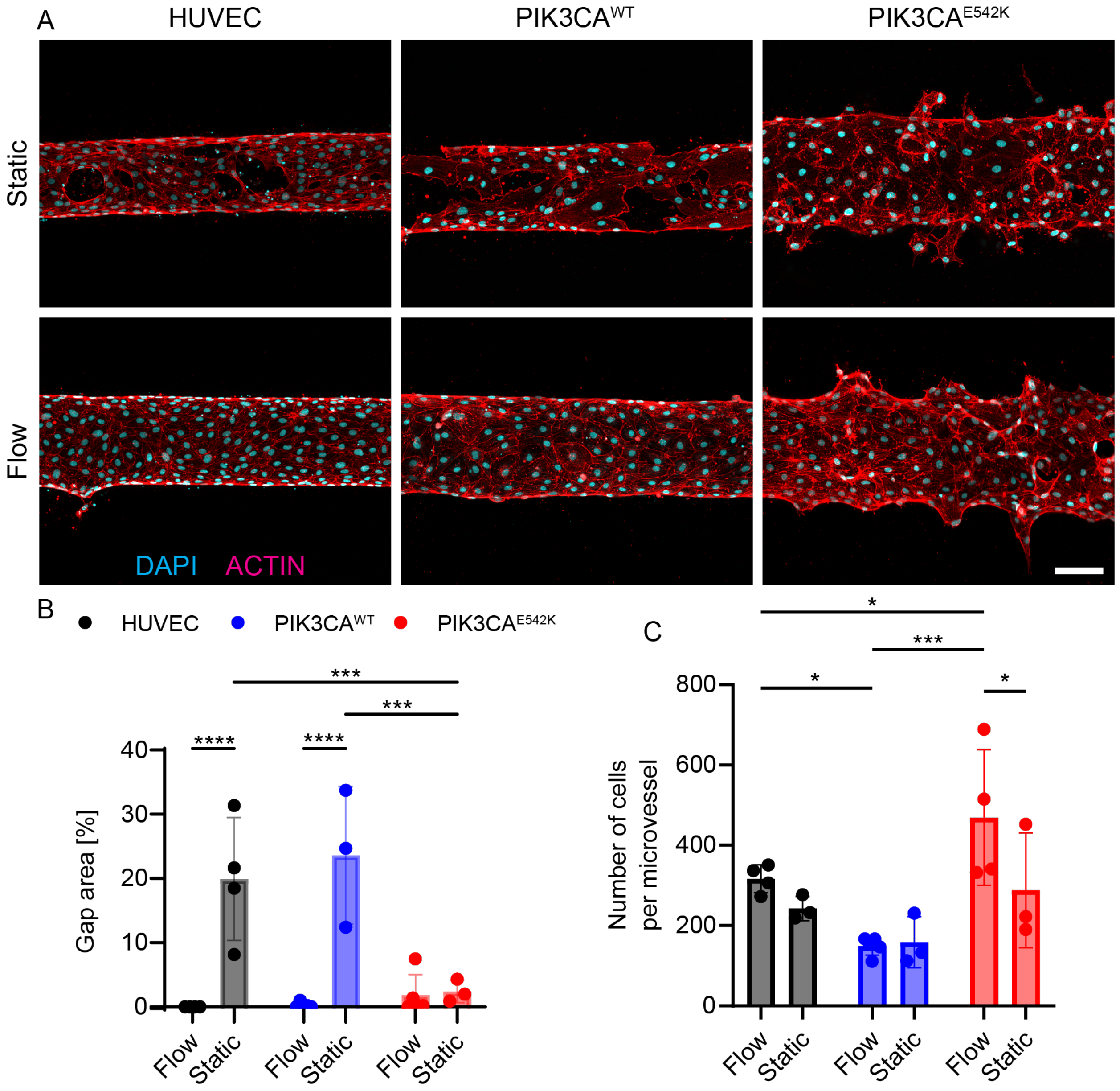


**Figure S5. Characterization of cellular coverage, sprouting and invasion in *PIK3CA$^{E542K}$* microvessels cultured under fully static condition** (**A**) Representative maximum intensity projection images of untransduced HUVECs, *PIK3CA$^{WT}$* or *PIK3CA$^{E542K}$* HUVECs cultured under fully static or continuous rocking for 48 hours. Microvessels were stained for DAPI (blue) and actin (red). Scale bar = 100 µm. (**B**) Quantification of percent gap area in control or mutant microvessels cultured under continuous flow or fully static condition (**C**) Quantification of total number of cells within each microvessels for the indicated genotype and culture condition.

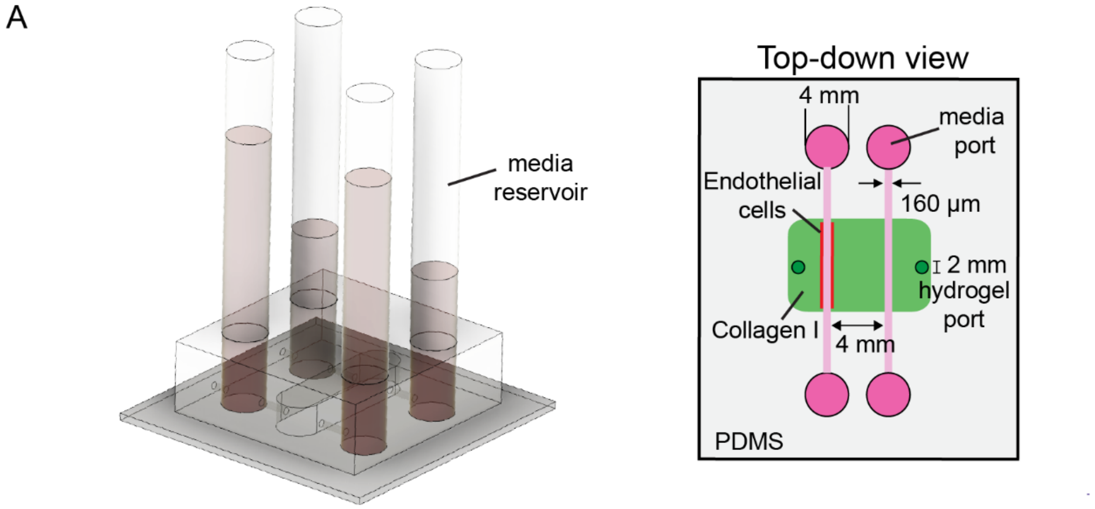


**Figure S6. Two-channel device used to establish transmural flow.** (**A**) Schematics of two-channel devices used to generate transmural flow across microvessels. Reservoirs are connected to the media ports and 20 mm $H_2O$ hydrostatic pressure gradient was applied to drive transmural flow into or out of the microvessels.

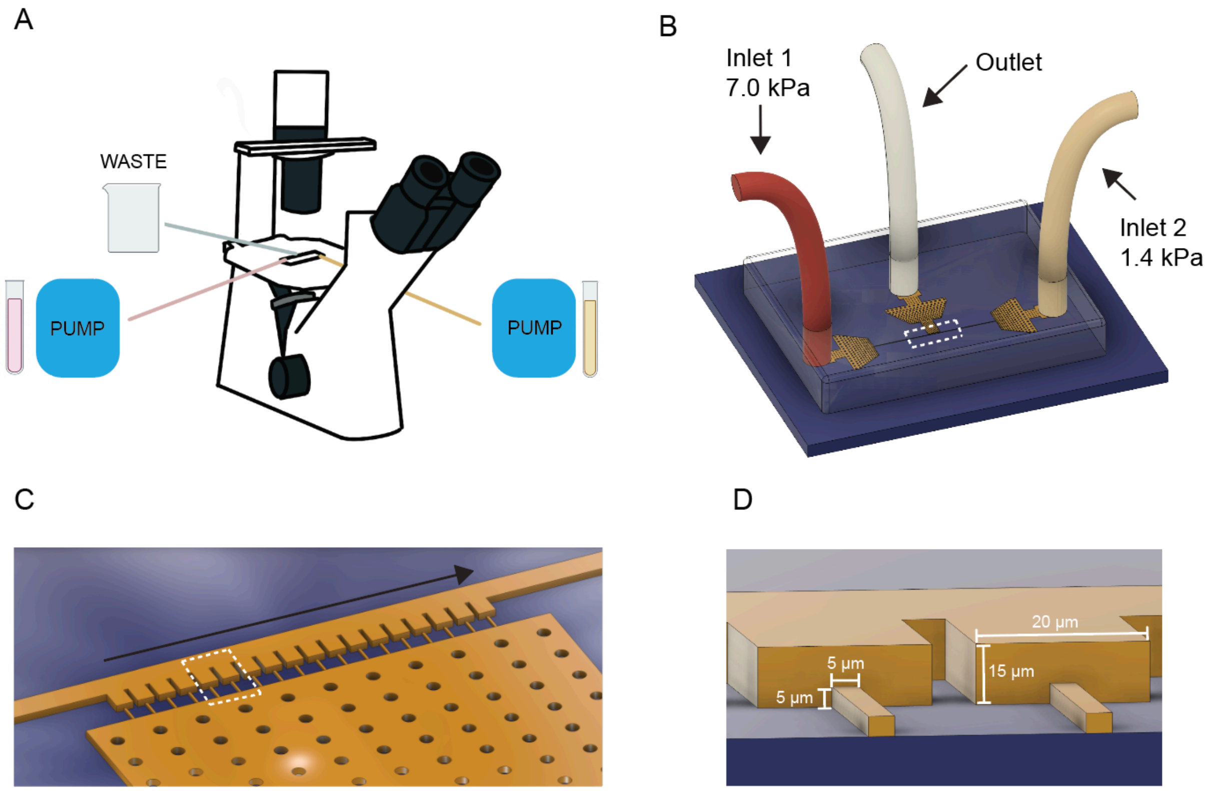


**Figure S7. Microfluidic micropipette aspiration setup and device parameters.** (**A**) Schematic of flow experiment setup and materials. Two constant-pressure pumps were connected to device inlets, pumping cell suspension (pink) and cell-free PBS (beige) into the device. All pumps and samples were kept on shelves at the same height as the microscope stage. (**B**) A 3D model of the chip on the microscope stage in (A), defining inlet pressures maintained by the pumps. (**C**) A 3D view of the dashed area in (B), showing the array of pockets that trap cells as they flow along the vector from Inlet 1 to Inlet 2. (**D**) A 3D view of the dashed area in (C), showing the dimensions of the pockets and the square constriction channel that the cells deform through under the applied pressure gradient.

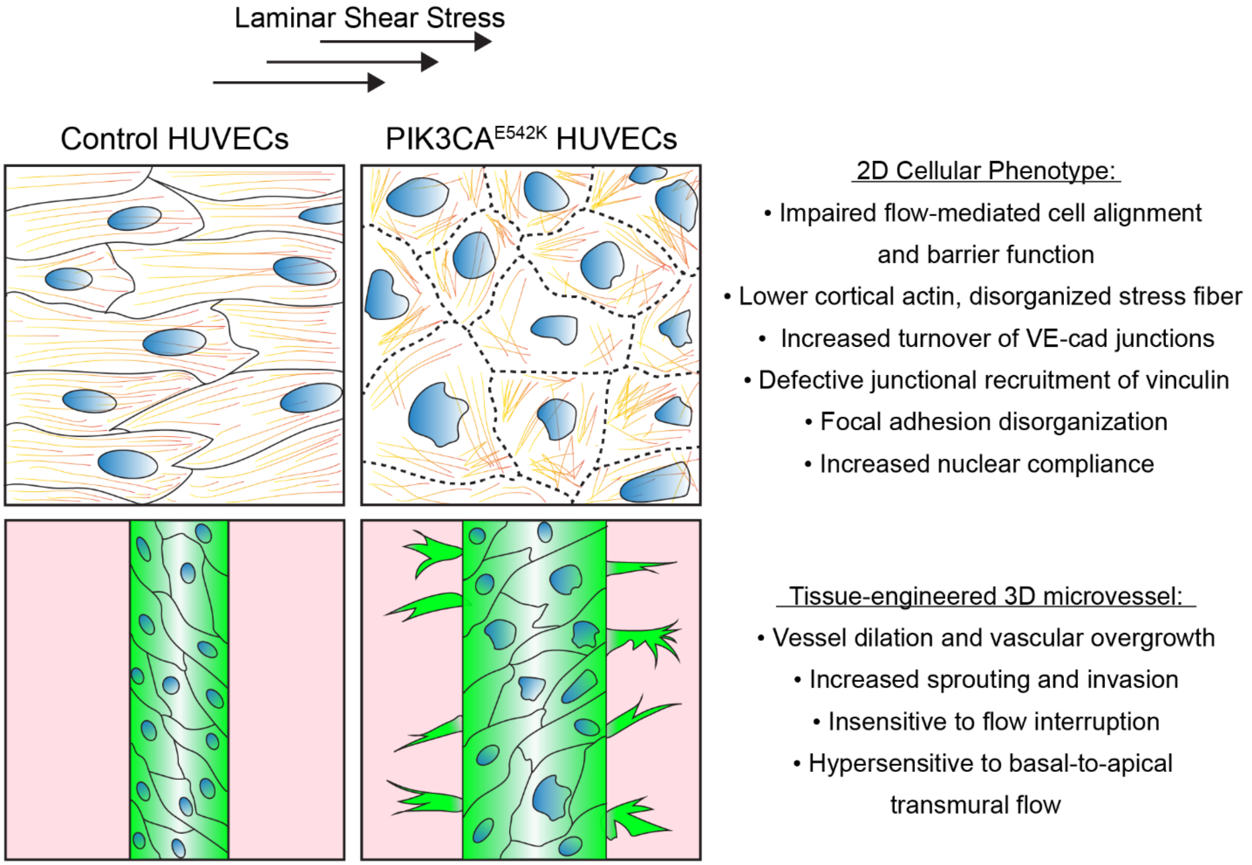


**Figure S8.** Schematic demonstrating cellular and vascular pathologies driven by the expression of *PIK3CA* activating mutation in HUVECs. Endothelial cells with *PIK3CA*$^{E542K}$ mutation fail to elongate, align, and establish barrier function under laminar shear stress. This is associated with increased nuclear compliance, defective cytoskeletal remodeling, increased junctional VE-cadherin turnover, and impaired recruitment of vinculin to cell-cell junctions and focal adhesion. Collectively, these results suggest that altered nuclear mechanosensing and mechanotransmission between nucleus and cytoskeleton contribute to the loss of flow response, vascular hyperplasia, and vessel sprouting and invasion in *PIK3CA*-driven vascular malformations.
